# Astrocytic μ-δ opioid receptor heterodimers mediate the antidepressant effects of ketamine’s metabolite

**DOI:** 10.64898/2026.05.31.727553

**Authors:** Shuo Yang, Ling-Jun Wang, Yunxiang Sun, Xiaoyan Ma, Yi Rong, Fong Tsz Hei, Tianxiang Li, Di Deng, Xiao-Xue Li, Zhaoxiang Zhang, Yan-Xia Liang, Xianzhang Bu, Tao Peng, Huan Xu, Chuang Wang, Xiang Cai, Qiang Zhou

## Abstract

A deeper understanding of the targets and mechanisms of fast-acting antidepressants, exemplified by ketamine, remains indispensable for better therapeutic strategies and understanding depression. Beyond the canonical neuron-centric NMDAR inhibition hypothesis, brain opioid system and glia-mediated processes are increasingly implicated in ketamine’s antidepressant efficacy, yet their precise contributions remain poorly understood. Here, we demonstrate that one major metabolite of ketamine, (2R,6R)-hydroxynorketamine (HNK), selectively targets μ-δ opioid receptor heterodimers (μ-δ-ORs) on astrocytes. By promoting the formation and/or stabilization of μ-δ-ORs, HNK engages Gs-coupled signaling, elevates intracellular cAMP, phosphorylates CREB (p-CREB) levels and Ca²⁺ dynamics in astrocytes, and consequently restores key astrocytic proteins and functions in depression models. Disrupting μ-δ-OR assembly or Gs signaling abolishes HNK-mediated antidepressant responses both in vitro and in vivo. Collectively, astrocytic opioid receptor heterodimers are critical to antidepressant responses and HNK may serve as a prototype compound for targeting astrocyte dysfunction across a wide range of brain disorders.

## Introduction

Major depressive disorder (MDD) imposes a substantial global burden, underscoring the urgent demand for therapeutics that combine rapid efficacy with improved safety profiles^1–5^. Ketamine, a prototypical rapid-acting antidepressant in treatment-resistant populations, has catalyzed efforts to develop next-generation agents that preserve therapeutic potency while minimizing adverse effects and abuseliability^6,7^. Systematic elucidation of ketamine’s targets and mechanism provides opportunities to identify novel pharmacological leads and to refine existing models of mood disorder pathophysiology.

Current perceived frameworks largely attribute ketamine’s antidepressant effects to neuronal NMDAR inhibition, with a predominant emphasis on the parent compound and neuronal contributions. However, accumulating clinical and preclinical evidence indicates that astrocytes undergo profound alterations in MDD, including dysregulated expression of core astrocytic proteins such as GLT-1, Kir4.1, and GFAP^8–14^. Given that astrocytes orchestrate brain homeostasis and provide structural and metabolic support to neurons, restoration of key astrocytic structure and functions is likely required for achieving better and fuller therapeutic efficacy. Consistent with this premise, multiple antidepressant drugs modulate astrocytic physiology, although the underlying molecular targets remain poorly defined ^15,16^. Notably, a single sub-anesthetic dose of ketamine induces rapid hypertrophy of astrocytic somata and processes, modulates GFAP and Kir4.1 expression, enhances brain-derived neurotrophic factor (BDNF) synthesis and release, and restores GLT-1 expression^17^.

At the circuit level, ketamine exerts rapid disinhibitory effects through NMDAR blockade on inhibitory GABAergic neurons, thereby increasing excitation in the brain, which is widely regarded as central to its antidepressant action^18–21^. Concurrently, NMDAR inhibition in excitatory neurons elevates BDNF signaling and potentiates synaptic transmission^7,22–24^. Beyond glutamatergic mechanisms, ketamine engages a broader receptor spectrum, including the endogenous opioid system, which has long been implicated in both the etiology and treatment of depression. Opioid receptors are abundantly expressed across key affective brain circuits, including the hippocampus and prefrontal cortex, and their dysregulation has been linked to multiple psychiatric conditions including depression^25–28^. A comprehensive picture on the contribution of opioid system to depression is still lacking, hindered by the inherent complexity and heterogeneity of both systems. Importantly, pharmacological blockade of opioid receptors abolishes ketamine’s antidepressant effects in both humans and rodent models, implicating a functional requirement for opioid signaling^29–34^. Recent findings further indicate that ketamine targets μ-ORs on medial prefrontal cortex somatostatin-expressing GABAergic neurons to initiate antidepressant responses^35^. In addition to canonical homodimer configuration, opioid receptors form heterodimer complexes, including μ-δ-ORs, which exhibit distinct signaling and pharmacological properties^36,37^. Limited evidence suggests that selective engagement of μ-δ-OR heterodimers confer antidepressant effects in preclinical models while exhibiting reduced addiction liability, thereby highlighting their therapeutic potential^36,38,39^.

Ketamine undergoes rapid and extensive hepatic metabolism, with an elimination half-life of 2 -4 hours, in humans ^40^, implicating its downstream metabolites as substantive contributors to its antidepressant effects^41^. Among these, (2R,6R)-hydroxynorketamine (HNK) has garnered particular attention due to its rapid and persistent antidepressant efficacy in preclinical models and it is apparently required for ketamine’s antidepressant effects^41^. Importantly, HNK does not inhibit NMDARs at therapeutically relevant concentrations^23,42,43^, and it lacks several hallmark adverse effects associated with ketamine, including dissociation and abuse potential^44^. Although HNK shares the main downstream signaling events with ketamine, its direct targets remain unidentified^7,23,42,43,45,46^. Whether HNK modulates astrocytic functions or engages OR systems within the context of antidepressant action is poorly understood.

In the present study, we employ multimodal experimental approaches including behavioral pharmacology, high-resolution imaging (split-GFP), structural modeling (AlphaFold3), electrophysiology, Ca²⁺ and cAMP imaging, western blotting, and immunostaining, to demonstrate, across scales from single-cell analyses to behavioral paradigms in mice, that HNK directly targets μ-δ-OR heterodimers on astrocytes. This interaction activates Gs-dependent signaling cascades, elevates intracellular cAMP level, and rapid elevation of key astrocytic proteins and restoration of certain key astrocyte functions. These signaling events culminate in the rapid restoration of astrocytic protein expression and functional capacity. Crucially, disruption of μ-δ-OR formation or downstream Gs signaling *in vivo* abolishes the antidepressant efficacy of HNK, thereby establishing a causal and mechanistic link between astrocytic OR heterodimers and rapid antidepressant action.

## Results

### Rapid and sustained antidepressant effects of HNK in mouse depression models

Since ketamine is used for treating treatment resistant depression, and we have shown previously that ketamine exhibits rapid and sustained antidepressant effects in a mouse model of treatment depression via two-weeks daily injection of stress hormone adrenocorticotropic hormone (ACTH)^47^, we first examined HNK effects in this ACTH model. By using two widely used depression-like behavioral measures, forced swim test (FST) and sucrose preference test (SPT), we demonstrated significant depression-like behaviors in adult male mice after, comparing to vehicle-injected mice (Figure 1A, 1B). Injection of HNK led to a reversal of both measures at 1 h and 24 h (Figure 1B), indicating an antidepressant effect. Similar depression-like phenotypes and their reversal were also observed in a more widely used depression model of chronic unpredicted mild stress (CUMS) (Figure 1C).

**Figure 1:**
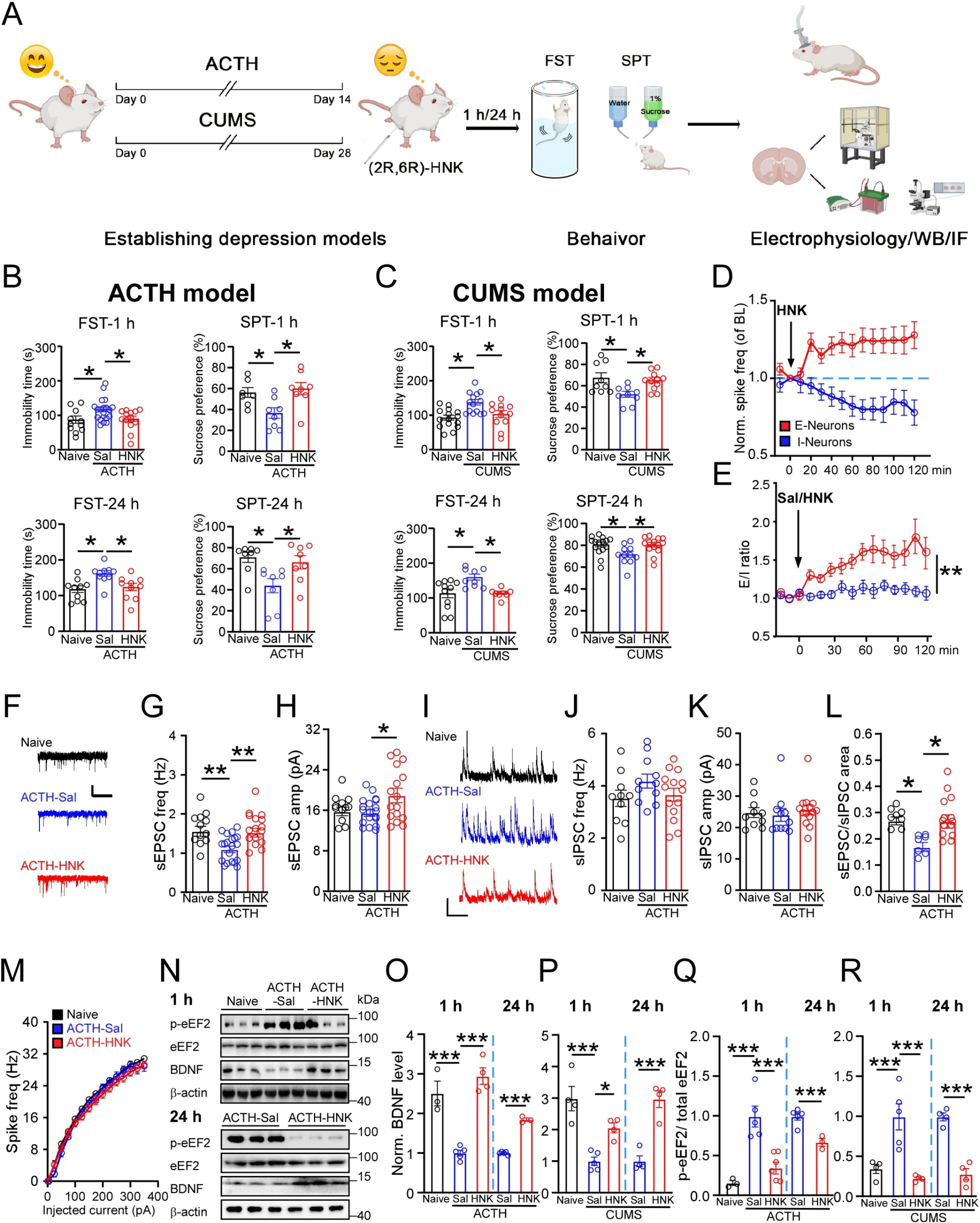
HNK elicits rapid and sustained antidepressant effects in mouse models of depression at the levels of behavior, brain activity, neuronal synaptic and biochemistry. (A) Schematic of experiments and procedures. ACTH and CUMS models were established, followed by behavioral tests (FST, SPT) and analysis. (B) Behavioral changes in ACTH-treated mice compared with naive and saline-injected controls after a single HNK injection (one-way ANOVA; n = 11, 20, 14, 7, 8, 8 mice for 1 h; n = 10, 10, 10, 8, 8, 8 mice for 24 h). (C) Behavioral changes in CUMS model mice compared with naïve mice, after a single HNK injection (one-way ANOVA; n = 15, 14, 12, 9, 9, 11 mice for 1 h; n = 11, 9, 8, 15, 11, 14 mice for 24 h). (D) *In vivo* spike rates of PFC E- and I-neurons in ACTH mice after HNK or saline injection (normalized to baseline (BL); arrow indicates injection). For HNK: E-neurons, 93 cells/10 mice; I-neurons, 23 cells/10 mice. For vehicle: E-neurons, 88 cells/8 mice; I-neurons, 21 cells/8 mice. (E) E/I ratio of spike rate before and after HNK injection in the same set of ACTH mice (one-way ANOVA; n = 10 mice per group). (F-H) Representative traces (F) and quantification of sEPSC frequency (G) and amplitude (H) in PFC E-neurons from naive mice, and ACTH mice injected with saline or HNK (one-way ANOVA; n = 12, 20, 16, 12, 19, 16 neurons/4 mice per group; scale bars, 10 pA/2 s). (I-K) Sample traces (I) and quantification of sIPSC frequency (J) and amplitude (K) in PFC E-neurons (one-way ANOVA; n = 10, 11, 14, 10, 10, 17, 8, 7, 15 neurons/4 mice per group; scale bars, 10 pA/2 s). L, Ratio of sEPSC/sIPSC areas in PFC E-neurons (one-way ANOVA; n = 8, 7, 14 neurons/4 mice per group). M, Intrinsic excitability of PFC E-neurons (n = 11, 16 neurons/4 mice per group). (N-R) Sample western blot images (N) and quantification of BDNF levels. All data normalized to mice injected with saline. BDNF: one-way ANOVA; n = 3, 5, 4, 4, 3 mice (ACTH); 4, 5, 4, 4, 4 mice (CUMS). p-eEF2/eEF2: one-way ANOVA; n = 3, 5, 6, 4, 3 mice (ACTH); 4, 5, 4, 4, 4 mice (CUMS). Data are mean ± s.e.m. *P < 0.05, **P < 0.01, ***P < 0.001.

Given the significance of the prefrontal cortex (PFC) in depression pathology and responsiveness to antidepressants^48–50^, we have focused on the PFC to elucidate the target of HNK. Our i*n vivo* recordings showed that HNK increased excitatory neuron (E-neuron) spiking and decreased inhibitory neuron (I-neuron) spiking (Figure 1D; Figure S1). This opposite change resulted in a rapid increase of E-I ratio measured using spike rate (Figure 1E), consistent with the well-established E/I increase with antidepressants including ketamine. In addition, E-neuron activity elevation occurred rapidly after HNK injection whereas I-neuron reduction developed gradually. The above distinct changes suggest that HNK likely direct effects E-neurons rather than via disinhibition.

Consistent with previous reports^51–53^, our whole-cell patch-clamp recording from identified PFC E-neurons further showed reduced spontaneous excitatory postsynaptic currents (sEPSCs) frequency and rapid elevation of both sEPSC frequency and amplitude by HNK (Figure 1F – 1H). In contrast, no significant alterations were observed in GABAergic inputs to the same set of PFC neurons, in ACTH mice without or with HNK injection (Figure 1I – 1K). The significantly lower E/I ratio of synaptic inputs (ratio of areas of sEPSCs/sIPSCs) in the PFC E-neurons from ACTH mice and its reversal by HNK suggests restoration of E/I balance (Figure 1L) ^54^. Since the intrinsic excitability of E-neurons was unaffected by ACTH or HNK injection (Figure 1M), the above findings indicate that the enhanced excitatory transmission in PFC is a main process associated with HNK’s rapid effects^55^. Consistent with prior studies, ACTH (Figure 1N - 1P) and CUMS mice (Figure 1P, 1R) exhibited significantly lower BDNF levels and higher p-eEF2/eEF2 ratios in the PFC, and both alterations were rapidly and persistently normalized by HNK (Figure 1N - 1R)^41,56,57^.

### HNK rapidly revers major changes in astrocytes in depression model mice

Beside neuronal alterations, significant alterations in the structures and functions of astrocytes have been demonstrated in both human depression patients and animal models^8–11,58^. Consistently, ACTH and CUMS mice showed reduced glial fibrillary acidic protein (GFAP) levels, a key marker for astrocytes, which were restored by HNK (Figure 2A)^59,60^. This reduction in GFAP level may indicate a reduced astrocyte density, and thus we used another astrocyte marker, S100β which labels cell body and partial processes of astrocytes. ACTH model mice showed lower S100β⁺ cell density and S100β level (Figure 2B, 2C), consistent with reduced astrocytes. A highly expressed and key astrocyte protein is Kir4.1 which regulates extracellular K^+^ concentration and neuron and astrocyte resting membrane potential and excitability ^61,62^. The level of Kir4.1 is altered in depression patients and models ^63,64^. Significantly lower PFC Kir4.1 level and its rapid reversal by HNK (Figure 2C).

**Figure 2:**
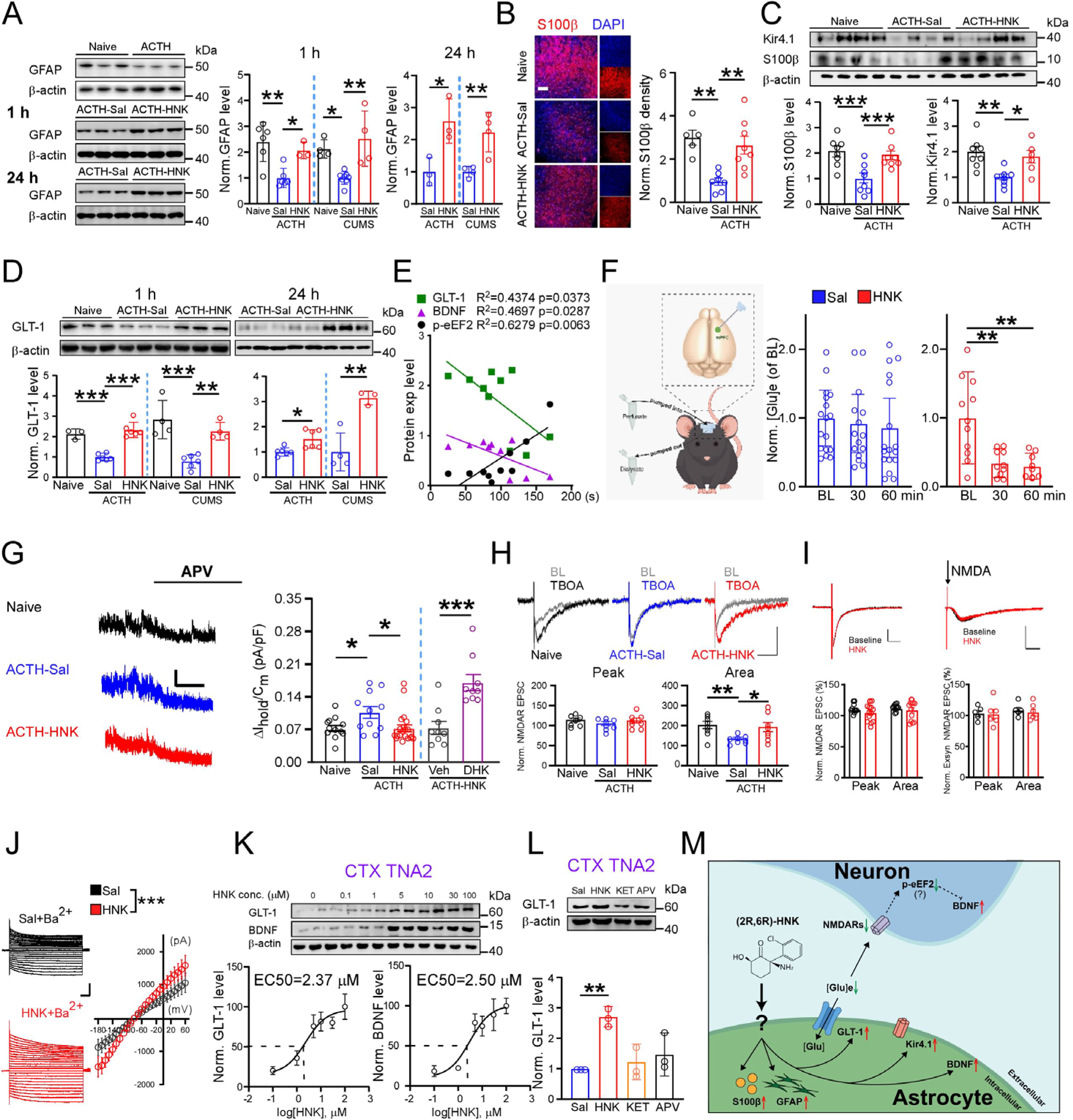
HNK restores astrocyte key protein expression and functions in depression model mice. (A) Sample images (left) and quantification of GFAP levels in depression models and its rapid (middle) and persistent (right) normalization after a single HNK injection (one-way ANOVA; n = 6, 6, 3, 4, 8, 4 mice for 1 h; n = 3, 3, 4, 4 mice for 24 h). (B) Sample images (left) and quantification of S100β+ cell density in PFC sections from naive, ACTH-Sal and ACTH-HNK mice (one-way ANOVA; n = 5, 7, 8 mice; scale bar, 200 μm). (C) Sample images (top) and quantification (bottom) of Kir4.1 and S100β levels in PFC (one-way ANOVA; n = 7, 8, 8 mice for S100 β; n = 8, 7, 6 mice for Kir4.1). (D) Sample images (top) and quantification (bottom) of GLT-1 levels in PFC (one-way ANOVA for 1 h, n = 3, 6, 6, 4, 6, 4 mice; two-tailed t-test for 24 h, n = 6, 6, 4, 3 mice). (E) Correlation analyses between PFC GLT-1, BDNF, p-eEF2/eEF2 levels and immobility time in FST within the same cohort (n = 10 mice). (F) Measurement of extracellular glutamate concentration ([Glu]e) in PFC using *in vivo* microdialysis before and after HNK or saline injection, normalized to pre-injection baseline (one-way ANOVA; n = 17, 11 mice). (G) Sample traces of ambient NMDAR responses in PFC E-neurons (left) and quantification (right). Higher ambient NMDAR response in ACTH-mice was reversed by HNK injection (one-way ANOVA; n= 12 neurons/5 mice, 11 neurons/5 mice, 20 neurons/5 mice, 9 neurons/5 mice, 9 neurons/5 mice; Scale bars, 15 pA/2.5 s). (H) Sample traces (top) and quantification of synaptic NMDAR-EPSCs (bottom) during baseline and after bath perfusion of DL-TBOA onto PFC E-neurons (one-way ANOVA; n = 8 neurons/5 mice per group; scale bars, 20 pA/50 ms). (I) HNK on isolated NMDAR-EPSCs (left): sample traces (top) and population results (bottom). Scale bars, 20 pA/100 ms. HNK on the isolated NMDAR responses to puffing of NMDA (right): Sample traces (top) and population results (bottom). Arrow indicates puffing of NMDA. Scale bars, 30 pA/100 ms. (J) Effect of HNK on Ba²⁺-sensitive currents: Sample traces (left) and current-voltage relationship (right; two-way ANOVA; n = 10, 11 neurons/3 mice per group; scale bars, 2 nA/50 ms). (K) Dose-response of HNK on GLT-1 (left) and BDNF levels (right) in CTX TNA2 cells (n = 4 experiments). (L) GLT-1 protein levels in CTX TNA2 cells 1 h after treatment with HNK, ketamine (KET) or D, L-APV (APV) (two-tailed t-test; n = 3 experiments). (M) Schematic summary of major findings in this figure. Data are mean ± s.e.m. *P < 0.05, **P < 0.01, ***P < 0.001.

Astrocytes express the highest levels of glutamate transporter-1 (GLT-1) among all brain cell types^65,66^, and we observed a significant lower GLT-1 in the PFC of both ACTH and CUMS model mice (Figure 2D)^6,67–73^, and reversal by HNK (Figure 2D)^59,74–77^. Since no alterations in GLT-1 mRNA level was observed (Figure S2,3), altered protein level is likely the main cause for the elevated GLT-1 level induced by HNK^76^. GLT-1, BDNF, and p-eEF2 levels were found to strongly correlate with FST immobility time (Figure 2E), and there was also a significant correlation between GLT-1 and BDNF levels (Figure S2), suggesting these markers track both depression-like behavior and its reversal by HNK^22,78^.

A main function of GLT-1 is to uptake glutamate from the extracellular space and hence modulate the extracellular glutamate level ^79,80^. We determined whether elevated GLT-1 enhances glutamate uptake using PFC microdialysis and we found that HNK significantly reduced [Glu]e (Figure 2F). Supporting this, NMDA receptors operating at resting membrane potentials are activated by [Glu]e, and their responses are termed ambient NMDAR responses^81^. Significantly higher ambient NMDAR responses were observed in the mPFC neurons from ACTH mice, and similar responses were obtained in naïve mice and HNK-injected ACTH mice (Figure 2G). In addition, bath application of selective GLT-1 inhibitor dihydrokainic acid (DHK) elicited significantly larger increase in the HNK-treated neurons, confirming the contribution of GLT-1 to these NMDAR responses (Figure 2G). Previous studies showed that GLT-1 has a significant contribution to the area of evoked NMDAR responses by modulating glutamate uptake from synaptic cleft^82^, providing another assay of GLT-1 function. Bath-applied pan-inhibitor of glutamate transporters DL-threo-beta-benzyloxyaspartate (DL-TBOA) produced smaller effects on isolated evoked NMDAR-EPSCs in PFC E-neurons from ACTH mice, whereas responses in HNK-treated ACTH mice resembled naïve controls (Figure 2H)^82,83^. Put together, these results indicate impaired glutamate uptake capacity in ACTH-mice that is reversible by HNK.

Suppression of NMDAR responses, including ambient NMDAR responses, plays a crucial role in ketamine’s antidepressant effect ^24,84^. Thus, an alternative explanation to reduced [Glu]e causing lower NMDAR-responses that HNK directly inhibit NMDARs. Patch clamp recording from PFC E-neurons reported no significant effect of acutely bath applied HNK on either synaptic (Figure 2I, right; Figure S4) or extrasynaptic NMDAR responses (Figure 2I, left; Figure S4) in the same set of neurons, at a HNK concentration (10 μM) showing *in vivo* antidepressant effects ^23,41–43^. This result is consistent with the absence of rapid effect of HNK on spiking in the I-neurons as what would be expected for a NMDAR inhibitor^85–88^. Thus, contribution of NMDARs to HNK’s antidepressant affects, if any, is likely indirect^18,56,89,90^. As Kir4.1 channels are altered in depression models^63,91^, we used the Ba^2+^ sensitivity of Kir4.1 channels^61,62^ and recorded Ba²⁺-sensitive currents in PFC astrocytes. The recorded current showed typical I-V relationship and Ba^2+^ sensitivity, both supporting Kir4.1 channel as a main contributor. A significant elevation was observed in neurons from HNK-treated ACTH-mice, supporting an elevated Kir4.1 level we have observed (Figure 2J; Figure S5).

In astrocyte cell line CTX TNA2 (Rat Brain Type I Astrocytes), HNK increased GLT-1 and BDNF with EC₅₀ values of 2.37 μM and 2.50 μM, respectively (Figure 2K). Interestingly, selective NMDAR antagonist DL-2-Amino-5-phosphonopentanoic acid (DL-APV) did not elevate GLT-1 in CTX TNA2 cells (Figure 2L), consistent with low astrocytic NMDAR expression and HNK not inducing NMDAR blockade^92–94^, and pharmacologically enhancing GLT-1 function reverses depression-like changes induced by chronic stress^9,10^. A summary of the major findings in Figure 2 is provided in Figure 2M.

### HNK targets μ-δ-opioid heterodimers on astrocytes

What is the target of HNK on astrocytes? Clinical and preclinical evidence suggests opioid receptors, particularly μ- and δ-opioid receptors (μ-ORs, δ-ORs) in both depression pathophysiology and antidepressant mechanisms of ketamine and its metabolites^28–31,95–99^. In CTX TNA2 cells, PFC slices, and *in vivo*, the μ-OR agonist [D-Ala2, NMe-Phe4, Gly-ol5]-enkephalin (DAMGO) mimicked HNK’s effects, whereas the μ-OR antagonist D-Phe-Cys-Tyr-D-Trp-Orn-Thr-Pen-Thr-NH2 (CTOP) or naloxone blocked HNK’s effects (Figure 3A; Figure S6). Similar changes in BDNF levels were observed in CTX TNA2 cells (Figure S6). HNK did not significantly affect GLT-1, BDNF, and p-eEF2/eEF2 levels in isolated astrocytes from μ-OR KO mice (Oprm1 KO; Figure 3B). Surprisingly, HNK failed to significantly modulate intracellular cAMP levels in CHO cells expressing exogenous μ-ORs (Figure 3C). Another opioid receptor subtype implicated in depression is the δ-ORs^26,100^. The agonist [D-Pen(2), D-Pen(5)]-enkephalin (DPDPE) mimicked HNK’s effects, while the antagonist naltrindole blocked HNK’s effects in CTX TNA2 cells, PFC slices, and *in vivo* (Figure 3D; Figure S7), with similar patterns of BDNF and p-eEF2/eEF2 modulation in CTX TNA2 cells (Figure S7). HNK did not significantly alter intracellular cAMP in CHO cells expressing δ-ORs (Figure 3E).

**Figure 3.**
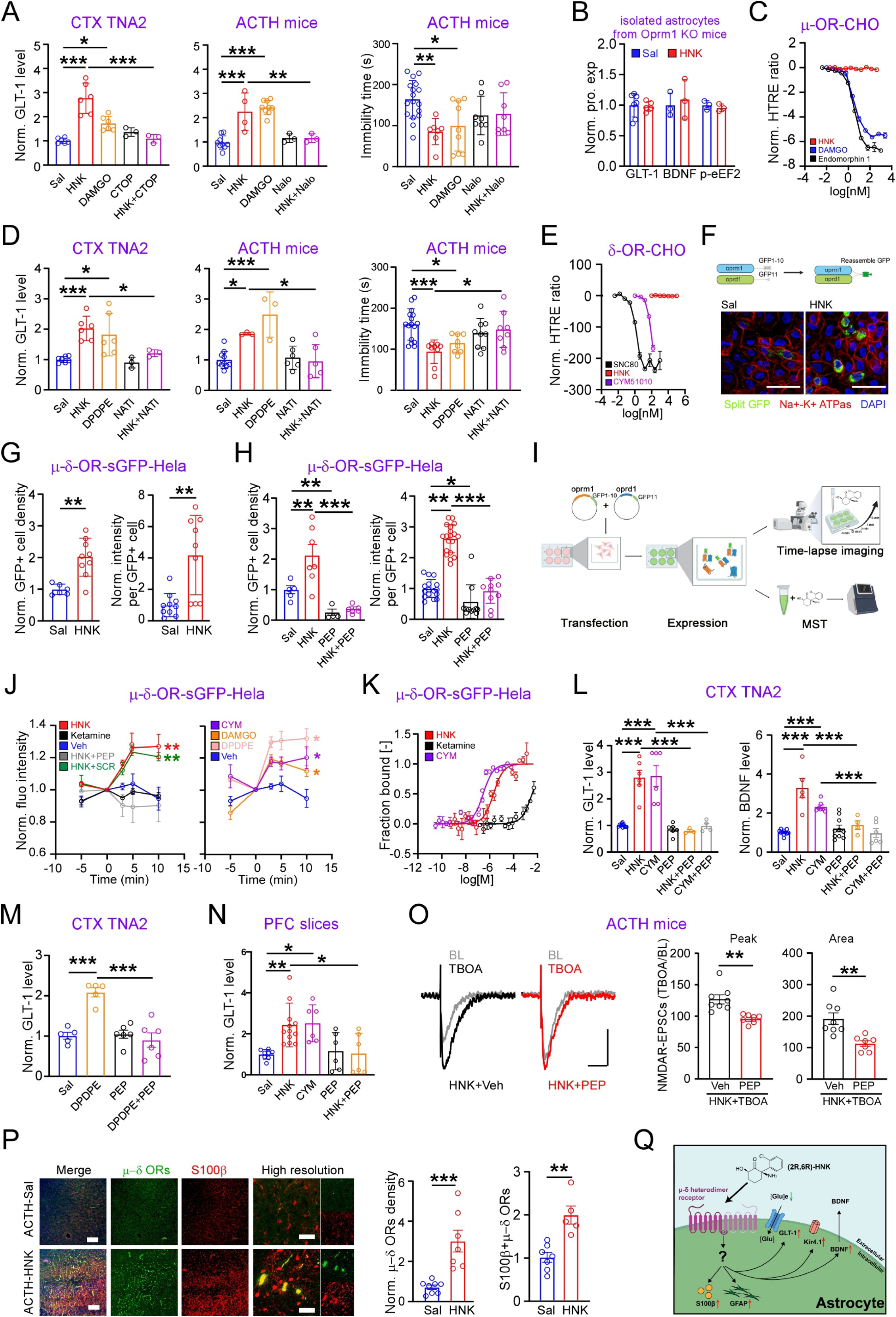
HNK targets μ-δ opioid receptors on the astrocytes. (A) Agonist of μ-opioid receptor DAMGO mimicked, while antagonists CTOP or Nalo blocked, the effects of HNK on GLT-1 levels in CTX TNA2 cells (left) and ACTH-mice (middle), and on FST immobility (right) (CTX TNA2, n = 3 - 6 experiments; *in vivo*, n = 9, 4, 9, 3, 3 mice; FST, n = 17, 7, 9, 8, 8 mice). (B) HNK on GLT-1 levels in astrocytes isolated from Oprm1knockout mice (n = 7, 5, 3, 3, 3, 3 mice). (C) HTRF analysis of cAMP levels in CHO cells expressing μ-ORs treated with indicated agents (10 concentrations, 2 replicates per compound). (D) Agonist of δ-opioid receptor DPDPE mimicked, while antagonist NATI blocked, HNK’s effects on GLT-1 levels in CTX TNA2 cells and ACTH-mice, and on FST immobility (CTX TNA2, n = 3 - 6 experiments; *in vivo*, n = 10, 3, 3, 6, 5 mice; FST, n = 16, 9, 8, 9, 8 mice). (E) HTRF analysis of cAMP levels in CHO cells expressing δ-ORs (10 concentrations, 2 replicates per compound). (F) Schematic of sGFP complementation assay for detecting μ-δ-OR heterodimerization (top), and sample images of GFP fluorescence in HeLa cells expressing μ-δ-OR-sGFP treated with HNK or vehicle, co-stained for Na⁺/K⁺-ATPase (cell membrane) and DAPI (nuclei; scale bar, 50 μm). (G) HNK on the density of GFP+ cells (left) and GFP fluorescence intensity per GFP+ cell (right; two-tailed t-test; n = 6, 9, 10, 9 cells/3 experiments). (H) PEP on HNK-induced changes in GFP⁺ cell density (left) and fluorescence intensity (right; one-way ANOVA; n = 5, 7, 4, 6 and 16, 19, 11, 10 cells/3 experiments). (I) Schematic of using sGFP in time-lapse imaging and MST experiments. (J) Time-lapse imaging of GFP fluorescence intensity in HeLa cells expressing μ-δ-OR-sGFP after incubation with indicated compounds (one-way ANOVA; n = 5 - 13 cells/3 experiments per compound). (K) HNK, ketamine and CYM on μ−δ-OR heterodimer from HeLa cells expressing μ-δ-OR-sGFP in MST assay (n = 4 experiments). (L) PEP pre-incubation on HNK- or CYM-induced elevated levels of GLT-1 (left) and BDNF (right), in CTX TNA2 cells (n = 6 experiments). (M) PEP pre-incubation on DPDPE-induced elevated GLT-1 levels in CTX TNA2 cells (n = 3 experiments). (N) GLT-1 levels in acute PFC slices treated with CYM, or HNK with PEP pretreatment (n = 6 - 11 slices/3 mice). (O) PEP pre-incubation on HNK’s effect on TBOA-induced changes in NMDAR-EPSCs in PFC E-neurons: sample traces (left) and quantifications (right; two-tailed t-test; n = 8 cells/4 mice, 7 cells/4 mice, 8, cells/4 mice, 7 cells/4 mice; scale bars, 20 pA/50 ms). (P) Immunostaining of μ-δ-ORs in PFC sections from ACTH-Sal and ACTH-HNK mice, co-stained with S100β, 1 h after HNK or saline injection: sample images (left) and quantifications (right; two-tailed t-test; n = 9, 7, 7,5 mice; scale bar, 200 μm/low resolution, 50 μm/high resolution). (Q) Schematic summary of the key findings in this figure. Data are mean ± s.e.m. *P < 0.05, **P < 0.01, ***P < 0.001.

The above findings suggest that HNK engages μ- and δ-opioid receptors in astrocytes through a mechanism independent of canonical cAMP signaling. One possibility is that HNK does not activate μ-OR or δ-OR homodimers, but engages μ–δ-OR heterodimers. These heterodimers are present in native tissues^101^, are formed in cells co-expressing both μ-ORs and δ-ORs, and they exhibit distinct pharmacological properties and functions^37,102–104^, and formation of μ-δ-OR heterodimers is observed with co-expression of both μ-ORs and δ-ORs^37,105^. To directly investigate potential interactions between HNK and μ-δ-OR heterodimers, we engineered μ- and δ-ORs fused to complementary split-GFP (sGFP) fragments, using GFP fluorescence to report heterodimer formation (Figure 3F; Figure S8)^106,107^. In HeLa cells expressing μ-δ-OR-sGFP, incubation with HNK significantly increased GFP intensity and the density of GFP-positive cells (Figure 3F, 3G), both consistent with a higher level of m-d-OR heterodimers in the presence of HNK, via either increased formation and/or stabilization. Prior incubation with a short TAT-fusion peptide (PEP) that disrupts the formation of m-d-ORs^108^ prevented the above increase (Figure 3H) with PEP alone also significantly reduced both measures, consistent with low level of spontaneous formation of m-d-OR heterodimers.

To determine the time course of HNK on the formation and/or stabilization of μ-δ-OR heterodimers, time-lapse imaging was used to monitor GFP fluorescence levels in the same set of cells before and after addition of HNK (Figure 3I). The increase in GFP fluorescence occurred within 2 min (first time point collected) of HNK application (Figure 3J), whereas no such increase occurred with peptide incubation, or following treatment with vehicle or ketamine (Figure 3J). The μ-δ-OR agonist CYM51010 (CYM)^38,109^ increased GFP fluorescence which was prevented by PEP preincubation (Figure 3J). Interestingly, increased GFP fluorescence was also induced by DEDPE or DAMGO (Figure 3J), consistent with previous findings that μ-δ-OR heterodimers can be activated by agonists and antagonists of μ-ORs and δ-ORs^36,37,110,111^. From this we infer the formation and/or stabilization of μ-δ-OR heterodimers are likely enhanced by HNK and other opioid receptor agonists.

To obtain a quantitative measure of the affinity of the above tested compounds on the μ- δ-ORs, we used MicroScale Thermophoresis (MST), on Hela cell lysates containing μ−δ-OR-heterodimers-sGFP incubated with HNK, and cell lysate containing GFP as a control (^112^, Methods; Sup. Figure S9). We have obtained a Kd value of 1.88 μM for HNK and 270.8 nM for CYM51010 on μ-δ-ORs (Figure 3K; ^38^). As a control, HNK did not show significant binding to GFP (Figure S9). Collectively, these findings support the notion that HNK binds to and either facilitates the formation of or stabilizes the formed μ-δOR heterodimers.

Preincubation with PEP blocked HNK or CYM-induced elevation of GLT-1 and BDNF in CTX TNA2 cells (Figure 3L, Figure S10), with similar disruption of DPDPE-induced effects (Figure 3M), indicating that activation of μ-δ-OR is critical to these subsequent effects. Similarly, incubation of PFC slices with HNK or CYM leads to elevated GLT-1, which was prevented by PEP preincubation (Figure 3N, Figure S11). In PFC E-neurons from ACTH-mice, PEP also blocked HNK-induced enhancement of glutamate transporter function/activity (Figure 3O). The above findings collectively suggest a high level of μ-δ-ORs after HNK treatment, which is further supported by PFC sections from ACTH-mice displaying higher densities of μ-δ-OR immunostaining and greater colocalization with the astrocyte marker S100β (Figure 3P). A summary of findings in this figure is presented in Figure 3Q.

### Activation of μ-δ-OR heterodimers activates Gs signaling and elevates intracellular cAMP level

The conventional signaling of μ-ORs and δ-ORs is mediated by activation of Gi/Gq signaling, reduced cAMP level and the associated signaling processes^113,114^. Activation of m-d-ORs has been shown to be insensitive to pertussis toxin^110,115^, and we thus first examined which G-protein signaling is engaged by HNK. PTX incubation did not affect elevation of GLT-1 in CTX N2A cells (Figure 4A). Inhibiting Gq signaling with FR900359 also had no significant effect (Figure 4B). We thus examined whether activation of Gs signaling may be involved. Viral Gs knockdown abolished the HNK-induced p-CREB elevation in HEK 293T cells (Figure 4C). Note that CYM-induced reduction in pCERB/CREB was still observed in cells expressing m-ORs, d-ORs or both, consistent with CYM’s lacking of selectivity over these ORs^38^. Similarly, preincubation with a selective Gs inhibitor NF449 blocked HNK-induced increases in p-CREB/CREB (Figure 4D) and GLT-1 (Figure 4E) in CTX TNA2 cells, with comparable effects in PFC slices from ACTH-mice (Figure 4F, 4G). NF449 also prevented HNK-induced enhancement of GLT-1 function in the PFC E-neurons (Figure 4H). These results implicate Gs signaling in downstream of μ-δ-OR activation by HNK.

**Figure 4:**
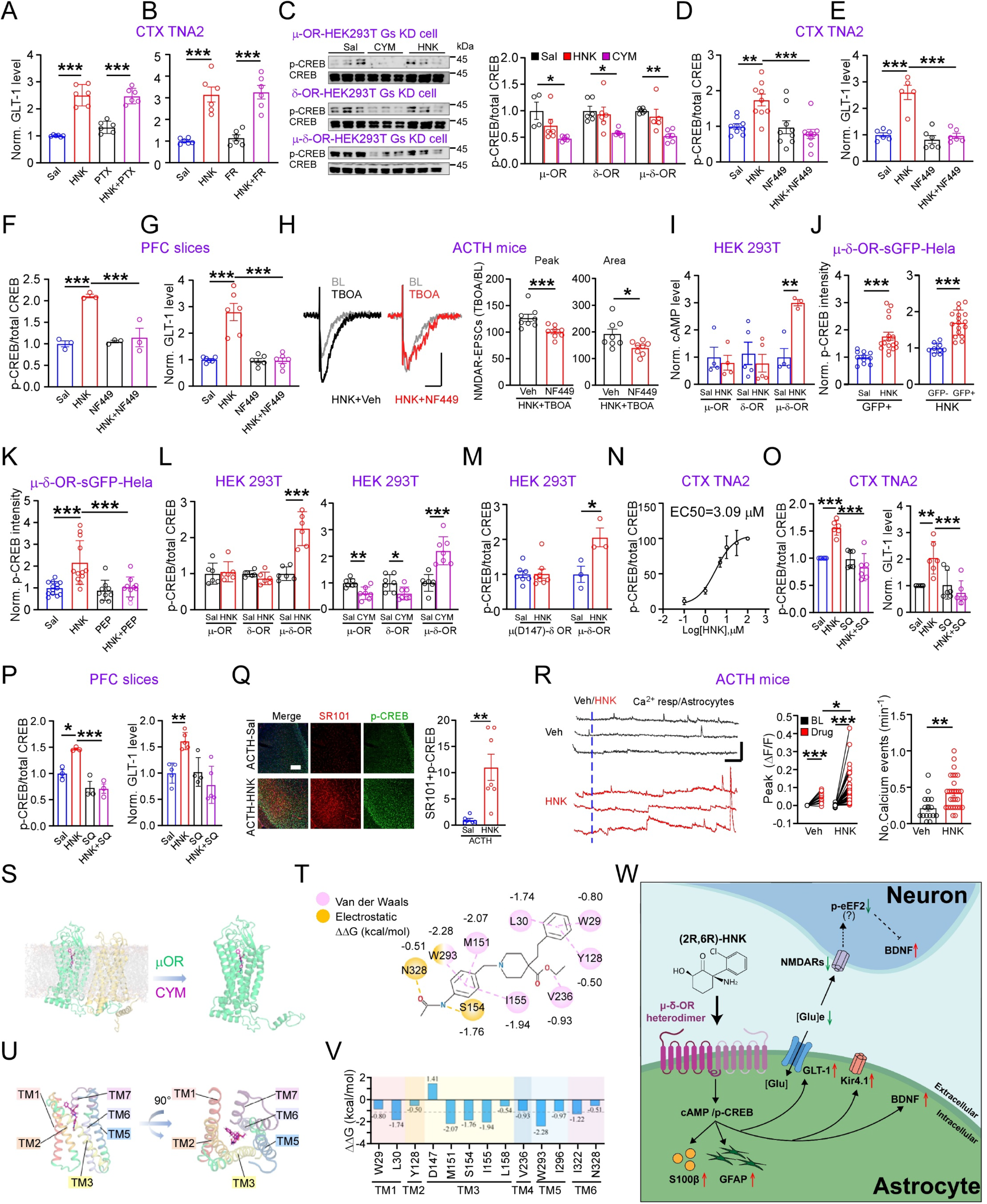
HNK-elicited intracellular signaling associated with activation of μ-δ opioid heterodimers. (A-B) PTX pre-incubation (A) or FR pre-incubation (B) on HNK-induced elevated GLT-1 levels in CTX TNA2 cells (n = 6 experiments each group). (C) Effect of Gs KD on HNK- and CYM-induced changes in p-CREB/CREB ratio in HEK 293T cells: sample images (left) and quantification (right) (one-way ANOVA; n = 6 experiments). (D-E) NF449 preincubation on HNK-induced changes in p-CREB/CREB (D) and GLT-1 (E) in CTX NA2 cells (one-way ANOVA; n = 9, 6 experiments). (F-G) NF449 pre-incubation on HNK-induced changes in GLT-1 levels (F) and p-CREB/CREB ratio (G) in acute PFC slices (one-way ANOVA; n = 6, 3 mice). (H) NF449 preincubation on HNK’s effect on TBOA-induced changes in NMDAR-EPSCs: sample traces (left) and quantification (right; two-tailed t-test; n = 8 cells/4 mice, each group; scale bars, 20 pA/50 ms). (I) HNK on cAMP level in HEK 293T cells (two-tailed t-test; n = 4 experiments). (J) p-CREB fluorescence intensity per sGFP⁺ HeLa cell (left), and in sGFP⁻ and sGFP⁺ cells treated with HNK (right; two-tailed t-test; n = 11, 17, 9, 17 cells/3 experiments). (K) PEP preincubation on HNK-induced elevated p-CREB fluorescence intensity in HeLa cells (one-way ANOVA; n = 13, 12, 9, 10 cells/3 experiments). (L) HNK and CYM on p-CREB/total CREB in HEK 293T cells (two-tailed t-test; n = 6, 7 experiments). (M) HNK on p-CREB/total CREB in HEK 293T cells (two-tailed t-test; n = 7 (μ(D149A)-δ-ORs), 3 (μ-δ-ORs) experiments). (N) Dose-response of HNK on p-CREB/CREB in CTX TNA2 cells (n = 4 experiments). (O) SQ preincubation on HNK-induced elevated p-CREB/CREB (left) and GLT-1 (right) levels in CTX TNA2 cells (one-way ANOVA; n = 6 experiments). (P) SQ preincubation on HNK-induced elevated p-CREB level (left), and GLT-1 level (right) in acute PFC slices (one-way ANOVA; n = 3 mice per group for p-CREB; n=5, 5, 4, 5 mice for GLT-1). (Q) Overlap between p-CREB+ and SR101+ cells in ACTH-mice: sample images (left) and quantification (right) (two-tailed t-test; n = 6, 7 mice; scale bars, 200 μm). (R) HNK’s effect on PFC astrocytic Ca²⁺ activity: representative traces (left; dotted line indicates addition of Veh/HNK; scale bars, 0.1 (ΔF/F)/60 s); peak Ca²⁺ responses (middle; two-tailed *t*-test; baseline vs. Veh, n = 16 cells/2 mice; baseline vs. HNK, n = 39 cells/3 mice; Veh vs. HNK); number of Ca²⁺ events per minute (right; two-tailed *t*-test; n = 18 cells/3 mice for Veh, 39 cells/2 mice for HNK). (S) Atomistic structure of CYM-OR heterodimer complex predicted by AlphaFold3. (T) Key interactions stabilizing CYM binding within the OR heterodimers. (U) CYM interaction profile across individual transmembrane helices of μ-OR within the heterodimer. (V) Per-residue binding free energy contributions to CYM binding. (W) Schematic summary of key signaling pathways identified. Data are mean ± s.e.m. *P < 0.05, **P < 0.01, ***P < 0.001.

Since Gs signaling is known to be associated with an elevated intracellular cAMP level, we measured HNK-induced cAMP levels in the HEK 293T cells expressing μ-ORs, δ-ORs, or both. Significantly higher cAMP levels were observed only in cells expressing both μ-ORs and δ-ORs (Figure 4I). Elevated intracellular cAMP level may increase p-CREB level^116^, and consistently higher p-CREB fluorescence intensity was observed in sGFP+ HeLa cells treated with HNK (Figure 4J), and significantly higher p-CREB intensity was observed in GFP+ vs. GFP- cells after both were treated with HNK (Figure 4J). In addition, this elevation was blocked by PEP pretreatment (Figure 4K; Figure S12A). A higher p-CREB/CREB ratio was also observed in HEK 293T cells expressing both μ-ORs and δ-ORs but not in those expressing only μ-ORs or δ-ORs (Figure 4L; Figure S12B). In contrast, CYM led to a higher p-CREB/CREB in HEK cells expressing both μ-ORs and δ-ORs but lower p-CREB/CREB in those expressing only μ-ORs or δ-ORs (Figure 4l; Figure S12C), consistent with CYM being an agonist of μ-ORs and δ-OR homodimers^38,117^. In contrast to HNK, ketamine did not alter p-CREB/CREB in HEK cells expressing μ-ORs and δ-ORs, but significantly reduced this ratio in cells expressing only μ-ORs (Figure S12D), consistent with ketamine primarily engaging μ-OR signaling^32,118^. We have recently identified D147A and Y148 in the μ-ORs as two key amino acids for binding of HNK to m-d-OR heterodimers, based on a-fold-3 analysis of the binding pocket and mutation of these 2 residues abolished HNK’s binding to μ-δ-OR heterodimers and loss of HNK’s antidepressant effects ^119^. Expressing μ-ORs with a D147A mutation abolished the elevation of p-CREB/CREB in HEK 293T cells that co-expressed μ-ORs and δ-ORs (Figure 4M; Figure S12E). The EC_50_ of HNK on elevating p-CREB/CREB was 3.09 μM (Figure 4N), consistent with other measures (Figure 2K, 3K).

The intracellular adenylate cyclase activity strongly modulates cAMP level. Consistently, artificially elevating cAMP level using dbcAMP dose-dependently elevated GLT-1 level in CTX TNA2 cells (Figure S13). In addition, adenylate cyclase inhibitor SQ22536 (SQ) blocked HNK-induced elevated p-CREB/CREB, GLT-1 levels in CTX TNA2 cells (Figure 4O, Figure S13), further suggesting that elevated cAMP is required for HNK’s effects. A similar effect of SQ was observed in PFC slices from ACTH mice (Figure 4P, Figure S13). To examine whether the HNK-induced elevation of p-CREB occurs in the astrocytes, we stained astrocytes using a marker SR-101 in live tissues^120^. A significantly higher percentage of SR-101^+^ cells were positive for p-CREB immunostaining in ACTH-mice injected with HNK (Figure 4Q). It is well established that elevated cAMP level is associated with higher frequency of spontaneous Ca^2+^ events in the astrocytes^121,122^. Live imaging in acute brain slices showed Ca^2+^ events occurred with a greater frequency and larger amplitude in HNK-treated ones (Figure 4R), consistent with an elevated cAMP/p-CREB^123^.

### Identification of CYM binding pocket in μ-δ-ORs using α-fold-3

Based on our established method and results on identifying the HNK binding pocket in μ-δ-OR heterodimers ^119^, we further examined whether the μ-δ-OR agonist CYM may bind to this region. Structural behavior of the μ-δ-OR heterodimer predicted by AlphaFold3 and evaluated by 400 ns of all-atom molecular dynamics simulations (Figure 4S) showed that the heterodimer retained a stable transmembrane architecture throughout the trajectory, with backbone RMSD values for the full complex and individual protomers remaining below 0.5 nm, indicating minimal structural drift (Figure S14). Despite the close structural similarity between μ-OR and δ-OR, CYM selectively binds to μ-OR. This preference arises from steric occlusion of the δ-OR orthosteric pocket by the μ-OR N-terminal domain, highlighting how heterodimerization reshapes ligand accessibility and creates functional asymmetry between protomers. CYM binding was primarily stabilized by an extended network of hydrophobic and aromatic interactions across TM1 (W29 and L30), TM2 (Y128), TM5 (V236), TM6 (W293 and I296), and TM7 (I322 and N328), whereas TM4 contributed minimally (Figure 4T). Consistent with this interaction pattern, CYM binding induced a broad perturbation across most transmembrane segments of the receptor, with the notable exception of TM4 (Figure 4U). Binding free energy decomposition (Figure 4V) further identified TM3 of the μ-opioid receptor as the primary interaction hub, where residues M151, S154, I155, and L158 contributed substantially to the binding affinity. Additional residues—including W29 and L30 (TM1), Y128 (TM2), V236 (TM5), W293 and I296 (TM6), and I322 and N328 (TM7)—also showed favorable interactions with CYM, suggesting that ligand binding is stabilized through a combination of deep-pocket anchoring within TM3 and peripheral hydrophobic and aromatic packing across surrounding helices. Collectively, these findings support the notion that CYM binds to and facilitates the formation and/or stabilization of the formed m-d-OR heterodimers. Based on elevated cAMP (Figure S13B) and p-CREB leading to higher GLT-1 level in astrocytes and elevated BDNF levels (Figure S11A). Based on pCREB leading to higher GLT-1 level in astrocyte^124^ and may also elevate BDNF levels^125,126^, a brief summary of findings in this figure is provided in Figure 4W.

### In vivo HNK antidepressant effects require activation of μ-δ-ORs

Thus far, we have defined HNK’s targets and signaling *in vitro*, but whether these mechanisms occur *in vivo* and are required for its antidepressant effects remains unclear. Because direct assessment of μ-δ-OR engagement in astrocytes *in vivo* is not feasible, we used PEP to disrupt μ-δ-OR formation. Two-photon imaging of cAMP levels *in vivo* in PFC astrocytes from ACTH-mice showed that HNK injection led to higher amplitude and area of spontaneous cAMP events (^127^, Figure 5A), which were prevented by prior PEP injection into the brain ventricles. *In vivo* two-photon imaging of astrocytes Ca^2+^ events demonstrated that HNK or CYM similarly increased spontaneous Ca²⁺ event amplitude and area, and these increases were prevented by prior PEP injection (Figure 5B). Pretreatment of PEP *in vivo* also abolished HNK-induced increases in GLT-1 (Figure 5C), BDNF (Figure 5D) and p-eEF2/eEF2 (Figure 5E) in PFC (Figure S15). PEP also blocked the elevation in sEPSC frequency and amplitude in PFC E-neurons in ACTH mice injected with HNK *in vivo* (Figure 5F). CYM also enhanced sEPSC frequency in the PFC E-neurons (Figure 5F). Behaviorally, prior PEP injection eliminated HNK’s effects in FST and SPT, whereas CYM mimicked these effects of HNK (Figure 5G, H). A single injection of CYM also resulted in persistent effects on FST and SPT in 24 h post-injection (Figure 5I). We then examined the effect of inhibiting Gs signaling using NF449. Prior *in vivo* injection of NF449 blocked HNK’s effect on GLT-1, BDNF, and p-CREB, p-eEF2/eEF2 in ACTH mice (Figure 5J), and abolished HNK’s antidepressant effects in ACTH-mice measured using FST and SPT (Figure 5K). Taken together, the above findings indicate that HNK activates μ-δ-OR heterodimers and Gs signaling to produce antidepressant effects *in vivo* (Figure 5L).

**Figure 5:**
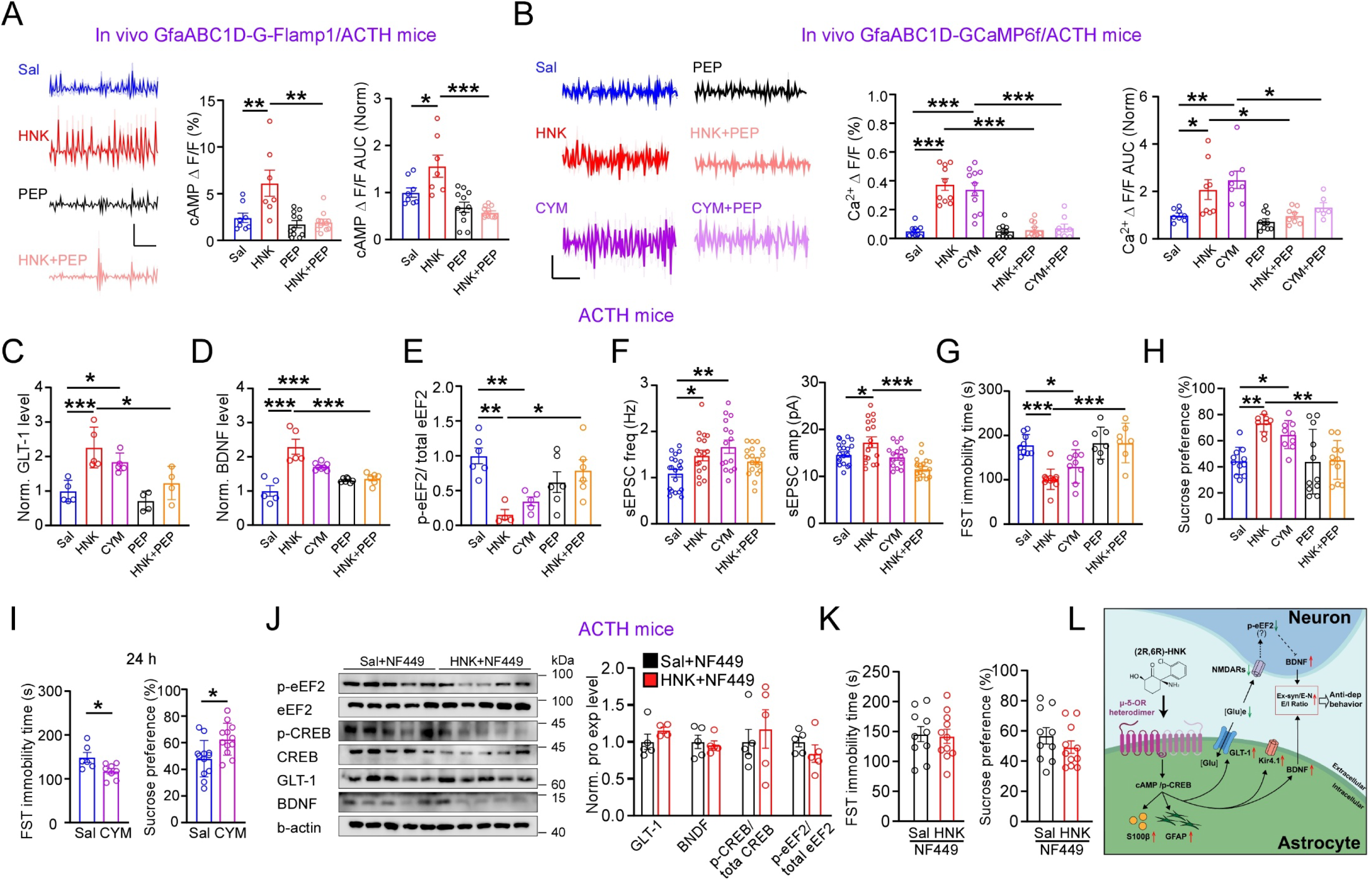
Activation of μ-δ opioid heterodimers mediates HNK’s anti-depressant effects. (A) Sample traces (left) and quantification of spontaneous *in vivo* cAMP events (frequency and AUC; right) in PFC astrocytes of ACTH-mice, 1 h after injection (one-way ANOVA; n = 8, 7, 11, 11 cells/4 mice per group; scale bars, 0.5 (ΔF/F)/50 s). (B) Sample traces (left) and quantification of spontaneous *in vivo* Ca²⁺ events in PFC astrocytes in ACTH-mice (right) (one-way ANOVA; for frequency, n = 10, 13, 12, 11, 11, 10 cells/4 mice; for AUC, n = 10, 10, 11, 10, 10, 8 cells; scale bars, 5(ΔF/F)/50 s). (C) HNK, CYM, or PEP pre-injection followed by HNK injection, on GLT-1 levels in ACTH-mice (one-way ANOVA; n = 5, 5, 5, 4, 4 mice per group). (D) HNK or CYM, or PEP pre-injection followed by HNK, on BDNF levels in ACTH-mice 1 h after CYM injection (two-tailed *t*-test; n = 5, 5, 7, 6, 6 mice per group). (E) HNK or CYM, or PEP pre-injection followed by HNK, on p-eEF2/eEF2 levels in ACTH-mice 1 h after CYM injection (two-tailed *t*-test; n = 6, 4, 5, 6, 6 mice per group). (F) HNK or CYM, or PEP pre-injection followed by HNK, on sEPSC frequency (left) and amplitude (right) in PFC E-neurons from ACTH-mice (one-way ANOVA; n = 19, 17, 15, 18 neurons/4 mice for frequency; n = 20, 17, 16, 18 neurons for amplitude/4 mice). (G-H) Effect of indicated treatments on FST (G) and SPT (H) in ACTH-mice (one-way ANOVA; n = 9, 11, 9, 7, 7 mice per group/FST; n = 11, 8, 9, 11, 11 mice per group/SPT). (I) Effect of a single injection of CYM on FST (left) and SPT (right) in ACTH mice 24 h after injection (two-tailed *t*-test; n = 6, 8 mice per group/FST; n = 12, 12 mice per group/SPT). (J) Intracerebral NF449 injection on HNK-induced changes in GLT-1, BDNF, p-CREB, and p-eEF2 levels in PFC of ACTH-mice: sample blots (left) and quantification (right; one-way ANOVA; n = 6 mice per group). (K) Effect of NF449 injection on HNK-induced changes in FST (left) and SPT (right) (two-tailed *t*-test; n = 10, 11 mice per group). (L) Summary of major findings in this study. Data are mean ± s.e.m. *, P < 0.05; **, P < 0.01; ***, P < 0.001.

## Discussion

In this study, we demonstrate that (2R,6R)-hydroxynorketamine (HNK) engages μ-δ-OR heterodimers on astrocytes, elevates intracellular cAMP, and initiates a signaling cascade that drives robust antidepressant effects *in vivo*. This signaling leads to rapid upregulation of astrocyte-enriched proteins, including GLT-1, GFAP and Kir4.1, together with increased BDNF levels. These molecular events are associated with functional restoration, encompassing enhanced glutamate clearance, improved potassium buffering, elevated excitation/inhibition (E/I) ratios at both synaptic and spiking levels, and potentiation of excitatory synaptic transmission, which are all hallmarks of antidepressant effects in animal models.

Integrating our current findings with the broader literature, we propose a dual-process framework underlying ketamine’s antidepressant action. The first component involves direct neuronal effects, wherein NMDAR inhibition and μ-OR activation converge to produce disinhibition and amplification of excitatory synaptic function, yielding a net increase in brain excitability and E/I ratio. The second component, delineated here, is mediated by HNK, which selectively targets astrocytic μ-δ-OR heterodimers. This astrocyte-initiated pathway exerts profound regulatory effects on neuronal architecture, synaptic plasticity, and global brain homeostasis. Consistent with this model, neither ketamine nor selective NMDAR antagonists activate μ-δ-ORs, and given ketamine’s relatively short half-life compared to HNK^46^, the metabolite likely constitutes a principal effector of ketamine’s antidepressant responses. This model is an expansion of Munguba^35^ to include astrocytes as a major player in ketamine’s antidepressant process.

### Key contribution of the opioid system to antidepressant effects

While functional expression of μ- and δ-ORs in astrocytes has been established^128–130^, the presence and physiological relevance of μ-δ-OR heterodimers in these cells have not previously been demonstrated. Importantly, the HNK-activated μ-δ-OR heterodimers engage Gs signaling, distinct from the canonical Gi/o coupling of μ-OR and δ-OR homodimers and from Gαz-mediated signaling reported for μ-δ-ORs^131^. This mechanistic distinction may provide a coherent explanation for the inconsistencies in prior studies on μ-OR and δ-OR contributions to antidepressant efficacy^29,31,38,132^, and reconciles apparently conflicting observations in ketamine pharmacological mechanisms^31,35,99,117,133^.

A central downstream effector is astrocyte-derived BDNF, which is robustly elevated by HNK and likely exerts strong influences on both astrocytes and neurons, thereby contributing critically to antidepressant outcomes. Elevated astrocytic cAMP is well established to promote cellular growth and maturation reminiscent of brain development^134^. The β-adrenergic receptors (β-ARs), a canonical Gs-coupled receptors in astrocytes, represent a similar and likely parallel signaling axis ^135,136^. Thus, determining whether β-ARs and μ-δ-OR heterodimers co-localize within the same astrocyte subpopulations is worthy of future exploration. Furthermore, the distribution and regulation of μ-δ-OR-expressing astrocytes across disease states, including depression and other brain disorders, remain to be systematically characterized^137^. Collectively, these findings reinforce the criticality of the endogenous opioid system in mood regulation and align with the high comorbidity between depressive disorders and opioid use/misuse^26,98,138–140^.

Upregulation of GLT-1 enhances extracellular glutamate clearance, thereby mitigating excitotoxic risk associated with increased neuronal activity. Concurrently, elevated cAMP and GLT-1 levels likely augment metabolic capacity, including glucose uptake and utilization, to support heightened energetic demands^141,142^. Increased intracellular cAMP and Ca²⁺ in astrocytes further modulate neuronal firing dynamics and cerebrovascular coupling, integrating metabolic and electrophysiological functions^143–147^.

We therefore propose that HNK orchestrates a coordinated astrocytic response that supports elevated neuronal function during recovery from depressive states. Notably, rapid GLT-1 upregulation reduces ambient glutamate levels and attenuates ambient/tonic NMDAR activation, which likely decreases p-eEF2/eEF2 ratios and facilitates neuronal BDNF upregulation^42,56^. This represents an additional astrocyte-to-neuron signaling axis mediated by extracellular glutamate modulation. While the present study focuses on acute functional restoration, whether HNK may also trigger longer-term astrocytic processes, including inflammatory modulation, remain to be elucidated.

### Targeting astrocytes to treat a spectrum of brain disorders

Given the extensive involvement of astrocytes in neurophysiology and their pervasive and excessive dysregulation across neurological and psychiatric disease setting, HNK-mediated restoration of astrocytic function may confer broad therapeutic benefits. Elevated intracellular cAMP levels in astrocytes serve as a master switch to promotes cellular maturation, metabolic support and homeostatic maintenance while restricting inflammatory and developmental features. More specifically, elevated cAMP triggers glycogenolysis and releasing energy substrates such as glucose; upregulates glucose transporters such as GLUT1, hence increasing the astrocyte’s capacity to import glucose from capillaries; increases GLT-1 expression, regulates AQP4 expression and permeability; restricts reactive astrogliosis and reduces the expression of pro-inflammatory cytokines^148^. These features, although not extensively examined here, represent critical targets for future investigation, particularly in disorders characterized by excessive reactive astrocyte responses, including Alzheimer’s disease, stroke, and traumatic brain injury^149,150^.

Astrocytic dysfunction, encompassing both structural alterations and dysregulated protein expression, is a convergent feature across numerous brain disorders, often mirroring alterations observed in depression. This convergence suggests that HNK may possess broad therapeutic applicability. For instance, GLT-1 downregulation is demonstrated in amyotrophic lateral sclerosis, Alzheimer’s disease, Parkinson’s disease, MDD, schizophrenia, ADHD, epilepsy, and traumatic brain injury, indicating that its restoration may confer widespread therapeutic effects^151–154^.

Altogether, the identification of μ-δ-OR heterodimers as a principal target of HNK carries substantial translational significance. Targeting these OR heterodimers has been associated with reduced tolerance and abuse liability comparing to μ-OR-selective agonists^38^, while their expanded signaling repertoire offers distinct pharmacodynamic advantages over OR homodimers^36,38^. Consistent with this profile, HNK exhibits minimal addictive potential^41,44,133,155^. Moreover, its apparent selectivity for μ-δ-OR heterodimers positions HNK as a prototype scaffold for the rational design of next-generation therapeutics that selectively target these receptor complex, enabling enhanced efficacy with reduced adverse effects across a spectrum of central nervous system disorders.

## Acknowledgments

We thank Timothy Mitchison for comments and suggestions, Zhou lab for discussion, and Oliver CF Zhou for editorial assistance.

## Author contributions

Conceptualization: Q.Z., X.C., C.W., S.Y.; Methodology: S.Y., LJ.W., YX.S., XY.M, Y.R., FT.H., TX.L., D.D., ZX.Z. XX.L.; Investigation: XZ.B., T.P., H.X.; Visualization: S.Y., Y.R., FT.H., YX.S.; Funding acquisition: Q.Z., C.X., YX.S.; Project administration: Q.Z.; Supervision: Q.Z.; Writing (original draft): S.Y., Q.Z.; Writing (review & editing): S.Y., Q.Z., X.C., C.W.

## Methods

### Animals and Drugs

This study was conducted under the guidance of management regulations for animal experiments and related activities at Peking University laboratory animal center of Shenzhen graduate school. All efforts were made to reduce the number of animals used and to minimize the suffering and pain of animals. Male mice (age postnatal day 8 to 10 weeks) were used in this study: wildtype (Balb/c and C57BL/6J) and Oprm1 KO (C57BL/6J genetic background). ACTH and CUMS modeling were used to induce depressive-like behaviors based on previous studies. Balb/c mice were injected with Adrenocorticotropic hormone (ACTH, 1mg/KG, Shanghai Yuanye, S25868) for 14 successive days. Chronic unpredictable mild stress (CUMS) model was established by subjecting C57 mice to various stressors^76^.

(2R, 6R)-HNK (10 mg/KG, Sigma-Aldrich, SML1873), naloxone (10 mg/KG, MCE, HY-17417A), Naltrindole (10 mg/KG, Sigma-Aldrich, N115), CYM51010 (10 mg/KG, MCE, HY-104006) were injected intraperitoneally. PEP (50 nM, QYAOBIO), DAMGO (10 μM, MCE, HY-P0210), DPDPE (1 μM, MCE, HY-P1334), NF449 (10 μM, MCE, HY-112461A) were injected into cerebral ventricle.

### Behavioural experiments

#### Forced Swimming Test

Mice were tested in a round glass tub (20 cm in diameter, 50 cm in height) containing water (20 cm deep) at 24 - 26 °C. After 2 min acclimatization, behavior was recorded and analyzed via an automated video tracking system (Ethovision XT 11.5, Noldus) in the following 4 min. The duration of immobility, defined as the time spent in a vertical position attempting to climb out, was quantified from the recorded videos using ForcedSwimScan™ 2.0 software (Clever Sys Inc).

### Sucrose Preference Test (SPT)

Mice were exposed to 1% sucrose solution and water for 24 h separately, and the sucrose and water bottles (left or right) were switched every 12 h, followed by water deprivation for 12 h before SPT. SPT procedure was performed for 1 h, the positions of bottles were changed after 30 min. Bottles were weighed before and after the test, and differences indicated the intake from each bottle. Sucrose preference was calculated as the percentage of sucrose intake to sum intake of water and sucrose.

### In vitro recordings and analysis

Mouse brains were rapidly removed and placed in an ice-cold cutting solution containing (in mM): 110 choline chloride, 7 MgSO_4_, 2.5 KCl, 1.25 NaH_2_PO_4_, 25 NaHCO_3_, 25 D-glucose, 11.6 sodium ascorbate, 3.1 sodium pyruvate, and 0.5 CaCl_2_ gassed with 95% O_2_ and 5% CO_2_. Coronal frontal sections (350 μm) were cut on a Leica VT1200S tissue slicer (Leica, German) in cutting solution.

Slices were transferred to a holding chamber with artificial cerebrospinal fluid ACSF containing (in mM): 127 NaCl, 2.5 KCl, 1.25 NaH_2_PO_4_, 25 NaHCO_3_, 25 D-glucose, 2 CaCl_2_, and 1 MgSO_4_, and allowed to recover for 30 min at 32 ℃, then kept at room temperature for at least 1 h before recording. Individual slices were transferred to the recording chamber on an Olympus microscope (BX51WI) with a ×40 water-immersion differential interference contrast objective. Slices were constantly perfused at room temperature (23–26 ℃) with oxygenated aCSF (4–5 mL/min). Recordings were made from layer 2/3 of the PFC, in a depth of about 50–100 μm in the slices. Data were acquired using HEKA EPC10 double patch clamp amplifier (HEKA). Signals were acquired at a sampling rate of 10 kHz and filtered at 2 kHz. Series resistance of the recording pipette was between 10 and 25 MΩ. Neurons with holding current > −200 pA (at −70 mV) were excluded from data analysis. Pyramidal neurons were identified by their apical dendrite and triangular somata.

To record spontaneous excitatory post-synaptic currents (sEPSCs), somatic whole-cell voltage clamp recordings (held at −70 mV) were obtained. Recording electrodes were filled with (in mM): 128 K^+^-gluconate, 10 NaCl, 2 MgCl_2_, 10 Hepes, 0.5 EGTA, 4 Na2ATP, and 0.4 NaGTP. To record spontaneous inhibitory post-synaptic currents (sIPSCs), somatic whole-cell voltage clamp recordings (+5 mV) were obtained. To isolate NMDAR EPSCs, neurons were held at −60 mV in the presence of NBQX (10 μM), picrotoxin (50 μM), Mg2+ (0.5 mM). For measuring the ambient NMDAR responses, slices were bathed in aCSF containing NBQX (10 μM), picrotoxin (50 μM), and Mg2+ (0.5 mM), with neurons held at +40 mV. D-APV (100 μM) was added into the recording chamber via a thin tube positioned at the edge of the recording chamber. Recording pipette solution contains (in mM): 125 CsMeSO_4_, 5 NaCl, 1.1 EGTA, 10 HEPES, 0.3 Na2GTP, 4 Mg-ATP, and 5 QX-314.

In experiments using TBOA, 10 μM HNK, 50 nM PEP, 10 μM NF449 were applied in the bath at least 1 h. NMDAR EPSCs were recorded in the presence of no TBOA (baseline) and 20 μM TBOA, in the same neurons. For measuring the ambient NMDAR responses with 10 μM HNK or (300 μM) DHK incubation (1 h), d-APV was fast perfused into the recording chamber by a tube positioned at the edge of the recording chamber.

### In vivo multi-channel electrode single unit recording

Mice were anesthetized with isoflurane (∼ 1% in a gas mixture) in a stereotaxic apparatus (RWD Life Science Co., China). Multi-wire electrode making and implantation were performed as described before^156^. After surgery, mice were allowed to recover for 7 - 14 days. During recording of spontaneous spiking, mouse could move freely in the recording box (390*180*180 mm), covered with clean bedding. Thirty min baseline was recorded before HNK injection (10 mg/KG, i.p) and 120 min after injection. Individually recorded neurons were sorted using Plexon Offline Sorter. Excitatory and inhibitory neurons classification, E/I ratio calculation were performed as described before^157^. Power spectrum densities and gamma oscillation were analyzed were analyzed using Neuro Explorer.

### Microdialysis

The monitoring device comprised of a double-lumen dialysis probe was inserted inside brain tissue. After implantation, a catheter was connected to a battery-operated pump that continuously perfused the brain with ACSF at rate 0.3 mL/min. The perfusion fluid was isotonic to the brain interstitial fluid to permit passive diffusion of small solutes from the interstitial fluid, and collected in small vials at hourly intervals. Level of glutamate was quantified.

### Cell culture

Cell lines used for imaging was Hela cell, and for western blot were CTX-TN2 (rat astrocyte) and HEK 293T (human, kidney). All cell lines were incubated in Dulbecco’s modified eagle medium (DMEM) supplemented with 10% fetal bovine serum (Gibco), 100 U/mL of penicillin, and 100 μg/mL of streptomycin (Gibco), and maintained at 37 ℃ with 5 % CO_2_ in incubator.

Plasmid transfection and Expression of μ-OR, δ-OR or μ-δ-OR-sGFP

Oprm1- ZsGreen plasmid encoding Mus musculus μ-OR (NM_001302793.1), oprd1-mcherry plasmid encoding Mus musculus δ-OR (NM_013622.3), oprm1-GFP1-11 plasmid encoding μ-OR::GFP1-10, oprd1-GFP11 plasmid encoding δ-OR::GFP11 were constructed from Hanbio Biotechnology company (China). The sequences of GFP1-10 and GFP11 were adapt from^158^. Plasmid were transfected using MegaTran 2.0 (Origene, China) into cultures of Hela or HEK 293T cells grown on a 6-well plate. Forty-eight hours after transfection, those cells were treated by drugs and live-imaged, fixed with 4 % paraformaldehyde for later imaging or western blot.

### Western blot

Samples from whole brain and brain slices were separated by 12% SDS-PAGE and transferred to PVDF filters (0.45 mm, Millipore). Membrane was blocked with 5% non-fat dry milk solution for 1 h and incubated overnight at 4 ℃ with rabbit polyclonal antibodies against p-eEF2 (1:1000, Cell signaling Technology, 2331), rabbit polyclonal antibodies against eEF2 (1:1000, Cell signaling Technology, 2332), rabbit polyclonal antibodies against GLT-1 (1:5000, Proteintech, 22515-1-AP), mouse monoclonal antibodies against GFAP (1:1000, Cell signaling Technology, 3670), rabbit monoclonal antibodies against BDNF (1:1000, Abcam, ab108319), rabbit polyclonal antibodies against β-actin (1:10000, Proteintech, 20536-1-AP), rabbit monoclonal Antibody against GAPDH(1:1000, Cell signaling Technology, 5174), rabbit polyclonal antibodies against p-CREB (1:1000, Proteintech, 28792-1-AP), mouse monoclonal antibodies against CREB (1:1000, Proteintech, 67927-1-Ig), rabbit polyclonal antibody against Kir4.1 (1:10000, Proteintech, 12503-1-AP), rabbit polyclonal antibody against S100β (1:1000, Proteintech, 15146-1-AP). After washing, membrane was incubated with peroxidase-linked anti-rabbit/mouse/goat IgG antibody (1:5000, Sigma), and blots were developed using electrochemiluminescence reagent (4A Biotech) and visualized using a Chemidoc MP gel imaging analyzer (BioRad).

### Immunohistochemistry

To examine change of astrocytes and μ-δ-ORs in the PFC of ACTH mice following HNK administration, mouse brains were fixed with 4% (vol/vol) paraformaldehyde (PFA), dehydrated with sucrose (30%) and embedded in optimal cutting temperature compound (OCT). Tissues were cut in 30 μm sagittal sections using a freezing-sliding Microtome (Leica, Germany). Cryosections were wished in phosphate buffer containing 0.5% Triton X-100 (PBST), and incubated with primary antibodies (anti-S100β, 1:500, Proteintech, 15146-1-AP; anti-p-CREB, 1:500, Proteintech, 28792-1-AP; anti-μ-δ-ORs, 1:500, Kerafa, EMS007) in blocking solution and incubated at 4°C overnight. Then primary antibodies were washed away and sections were incubated with secondary antibodies (Alexa Fluor 546 goat anti-rabbit IgG, Alexa Fluor 488 goat anti-mouse IgG,1:2000, A11035, Invitrogen, UK) for 1 h. Sections were then counterstained with DAPI for 10 min. For analysis and quantification of immunoreacted areas, sections were imaged using confocal microscopy (Nikon, Japan).

To examine the formation of μ-δ-OR-sGFP, HEK 293T or Hela cells transfected with oprm1-GFP1-10 and oprd1-GFP11 respectively or simultaneously were fixed in 4% paraformaldehyde in PBS for 15 min, permeabilized with 0.1% Triton X-100 in PBS. Next, cells were blocked with 10% goat serum albumin (BSA)/PBST for 1 h at room temperature, stained for primary antibodies (anti-Na+-K+-ATPase, 1:500, Abam, ab76020; anti-p-CREB, 1:500, Proteintech, 28792-1-AP) and Alexa Fluor 546 conjugated secondary antibodies.

### In vivo two-photon imaging

Genetically encoded cAMP sensor rAAV-GfaABC1D-G-Flamp1 viruses and Ca2+ indicators rAAV-GfaABC1D-GCaMP6f viruses were obtained from BrainVTA company (China). ACTH mice were anesthetized with isoflurane (induction 3%, maintenance 1.5%), secured in a stereotaxic frame and received bilateral infusion of AAV (200 nL/side) in PFC (1.98 mm anterior to Bregma; 0.25 mm lateral to midline; 1.00 mm ventral from the cortical surface). Mice were allowed a 4-week recovery period. To minimize stress from head restraint, awake mice were habituated to the imaging apparatus for approximately 15 min. PFC imaging was performed using a Bruker Ultima Investigator two-photon microscope equipped with a ×25 objective (NA 1.1) and controlled by Prairie View software. Green fluorescent indicators were excited with a 920 nm laser, and emitted fluorescence was collected through 490–560 nm filters at a sampling rate of 1–2 Hz. The fluorescence intensity of ROIs was extracted using ImageJ software. Background-subtracted fluorescence intensity was used to calculate ΔF/F.

### In vitro Calcium Imaging

Mice were injected with AAV2/9-GFaABC1D-GCaMP6s (5E+12vg/ml, 300 nl, bilateral) into the prefrontal cortex (PFC) 28 days prior to experiments. Acute 300 µm brain slices were incubated in HNK (10 µM) for 1 hour. Slices were placed in a recording chamber under a CMOS camera (401D, Tucsen, China). Imaging was performed using a mercury lamp (25% power, OLYMPUS) with a 500 ms exposure time and no interval between frames. After a 1 min baseline recording, either Veh or HNK (10 µM) was bath-applied, and imaging continued for 10 min.

### Microscale thermophoresis assay

Drugs or endogenous opioid peptide at different concentrations were mixed with HEK 293T cell lysate expressed μ-OR- ZsGreen, δ-OR-mcherry, μ-δ-OR-sGFP or GFP and incubated for 10 min at room temperature. The specimens were loaded and measured on monolith NT.115 instrument (Nano Temper Technologies, Germany). The dissociation constant (Kd) was fitted by the MO. Affinity Analysis software.

### Computational method

To probe the molecular basis of CYM recognition within the μ-δ-OR heterodimer assembly, we first generated a structural model of the heterodimer. Although experimental studies have reported the existence of μ-δ-OR heterodimers, the specific three-dimensional configuration used here was independently predicted with AlphaFold3^159^. The same framework also suggested a putative CYM binding pocket, which served as the starting point for subsequent atomistic simulations. The predicted transmembrane heterodimer was inserted into a lipid bilayer and solvated using the CHARMM-GUI preparation pipeline^160^.

All molecular dynamics (MD) simulations were performed with GROMACS 2020.3 (Spoel V. GROMACS 2020 Source Code; Zenodo), employing the Amber19SB force field for the receptor^161^. CYM parameters were derived using AmberTools with GAFF^162^, and ligand partial charges were calculated through RESP fitting based on Hartree–Fock electrostatic potentials ^163^. The system was solvated in TIP3P water and ionized with Na⁺ and Cl⁻ to approximate physiological ionic conditions (0.15 M).

Simulations were conducted at 310 K and 1 bar using a velocity-rescaling thermostat^164^and a Parrinello–Rahman barostat ^165^, respectively. Long-range electrostatics were treated with the particle mesh Ewald method ^166^, applying a 1.4-nm real-space cut-off that was also used for van der Waals interactions. Periodic boundary conditions were imposed in all dimensions. To quantify CYM–receptor interactions, we computed residue-level contributions to binding using the MM/PBSA formalism^167^, enabling a decomposition of energetics across the heterodimer interface.

### Statistics

Statistical significance was assessed using an unpaired t-test or ANOVA followed by Dunnett’s or Tukey’s tests. Normalized values were calculated as percentage change over the baseline. Data values are presented as mean ± SEM. Statistical significance was established as P < 0.05 (*), P < 0.01 (**), and P < 0.001 (***).

## Extended Data Figures

**Figure S1:**
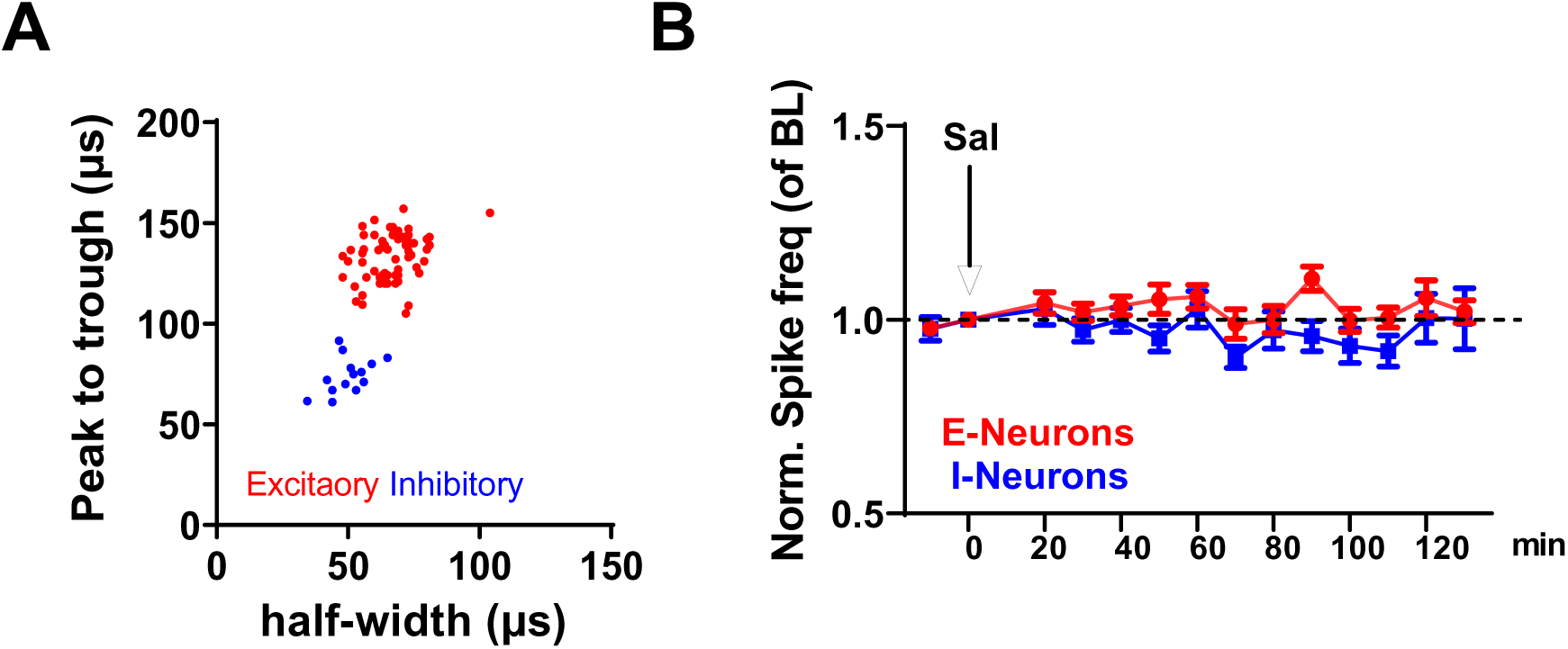
HNK’s impact on *in vivo* spike rate. (A) Criteria for identifying E- and I-neurons in PFC based on the peak to trough and half-width of their spike waveform. (B) Normalized spike rate (to pre-injection baseline) in E-neurons and I-neurons before and after vehicle/saline injection (arrow indicates injection; n = 88 E-neurons, 21 I-neurons/8 mice).

**Figure S2:**
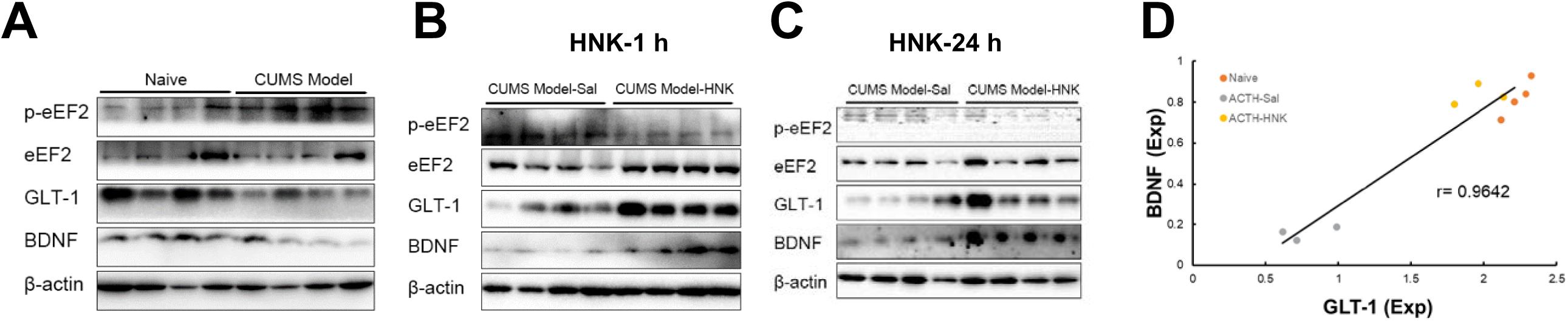
Western blot analysis of GLT-1, BDNF and p-eEF2/eEF2 levels in PFC from ACTH and CUMS model mice. (A-C) Sample western blot images from CUMS model mice (A), 1 h (B) and 24 h (C) after HNK or vehicle injection. (D), Relationship between BDNF and GLT-1 protein levels in PFC of naïve mice (orange), ACTH-saline mice (gray), and ACTH-HNK mice (yellow; n = 10 mice).

**Figure S3:**
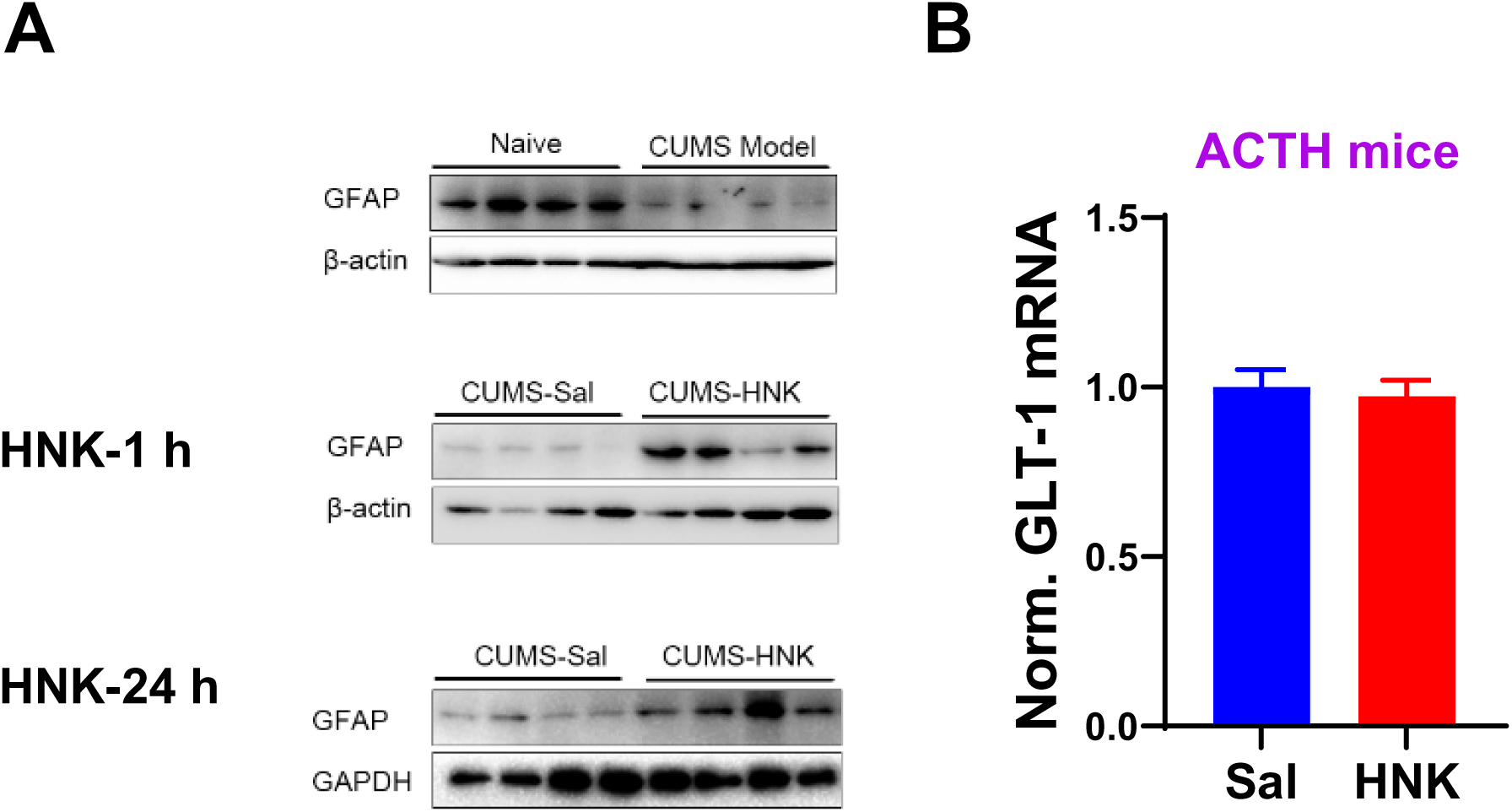
Restored GFAP levels in both ACTH and CUMS mouse models. (A) Sample WB images of GFAP in CUMS model mice 1 h or 24 h after HNK or vehicle injection. (B) GLT-1 mRNA levels in the PFC of ACTH-mice 1 h after HNK or saline injection (two-tailed t-test; n = 6 mice per group).

**Figure S4:**
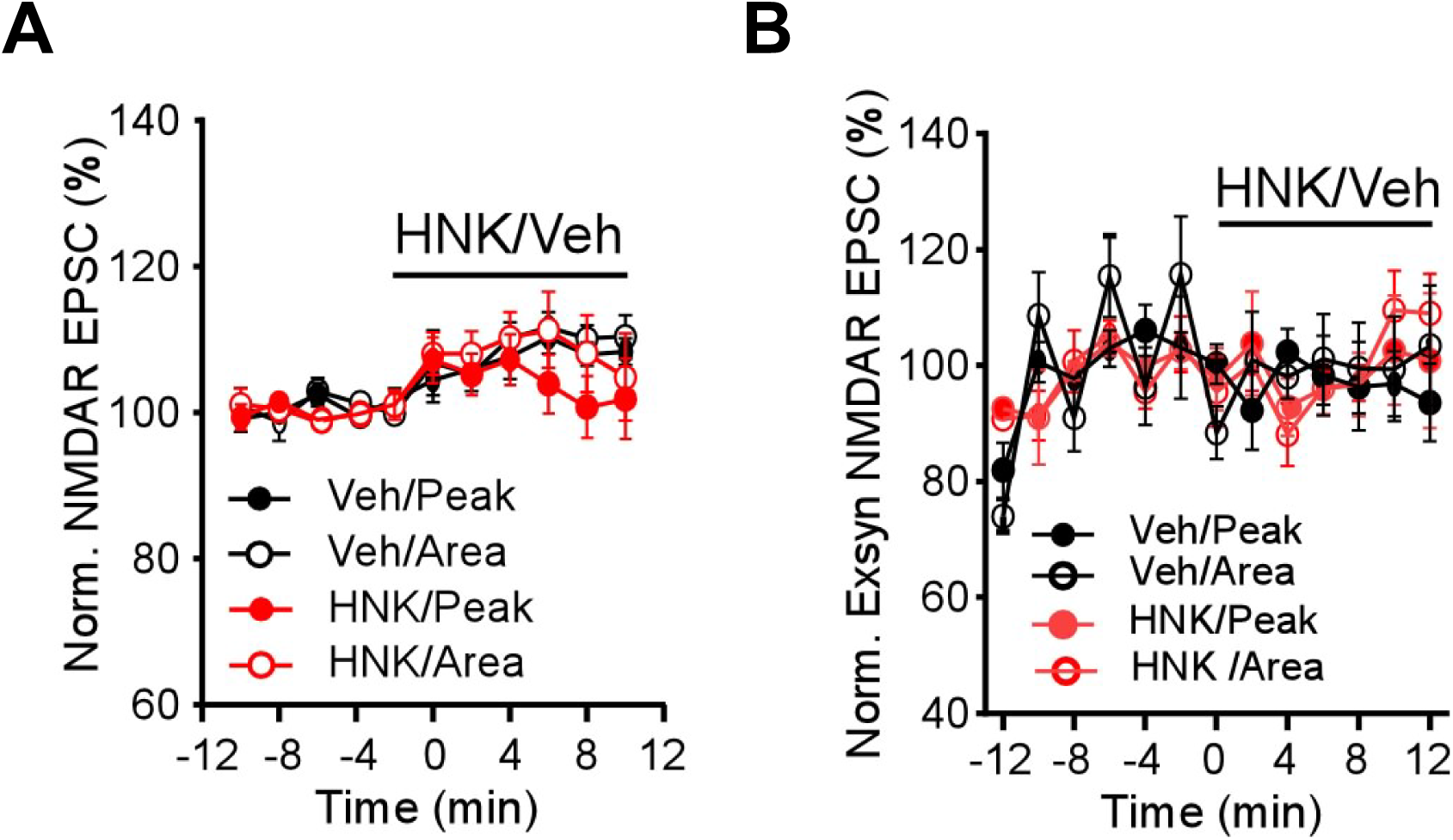
Time course of HNK on evoked NMDAR EPSCs or extrasynaptic NMDAR currents. (A) HNK on isolated NMDAR-EPSCs. Bar indicates HNK bath application. n= 11, 14 cell/4 mice. (B) HNK on the isolated NMDAR responses to puffing of NMDA. Bar indicates HNK bath application. n= 6 cell/3 mice per group.

**Figure S5:**
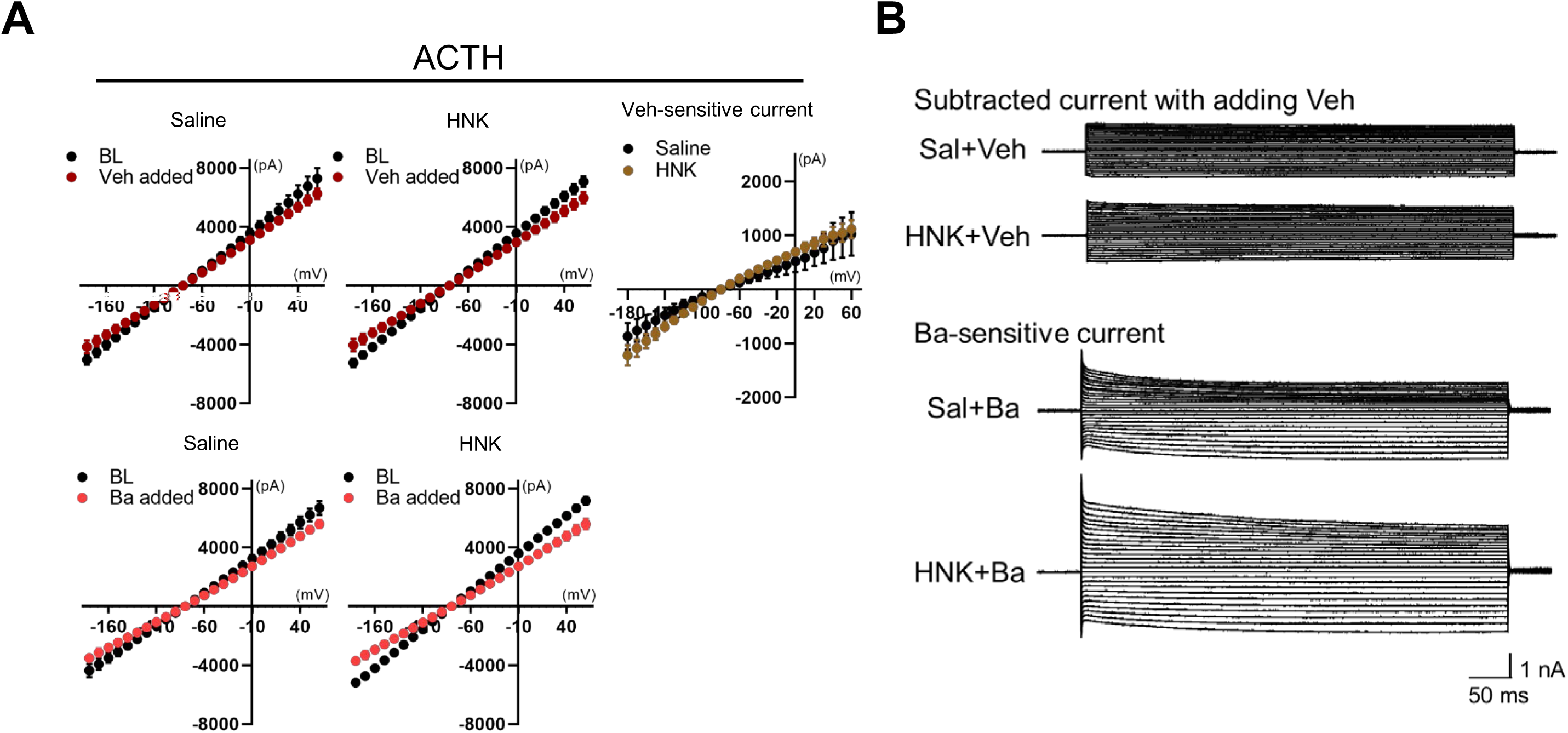
HNK increases Ba^2+^-sensitive current in astrocyte in ACTH-mice PFC slices. (A) Left and middle: I-V plots showing changes in currents after Veh (upper, n = 4 cells) or Ba^2+^ (lower, n = 10 cells) addition. Brain slices were pre-incubated with saline (left) or HNK (middle). Right: I-V plot showing the current obtained by subtracting the baseline current from currents recorded after Veh was added. (B) Upper: Representative current traces elicited by addition of Veh. Lower: Representative Ba²⁺-sensitive currents (scale bar, 1 nA/50 ms).

**Figure S6:**
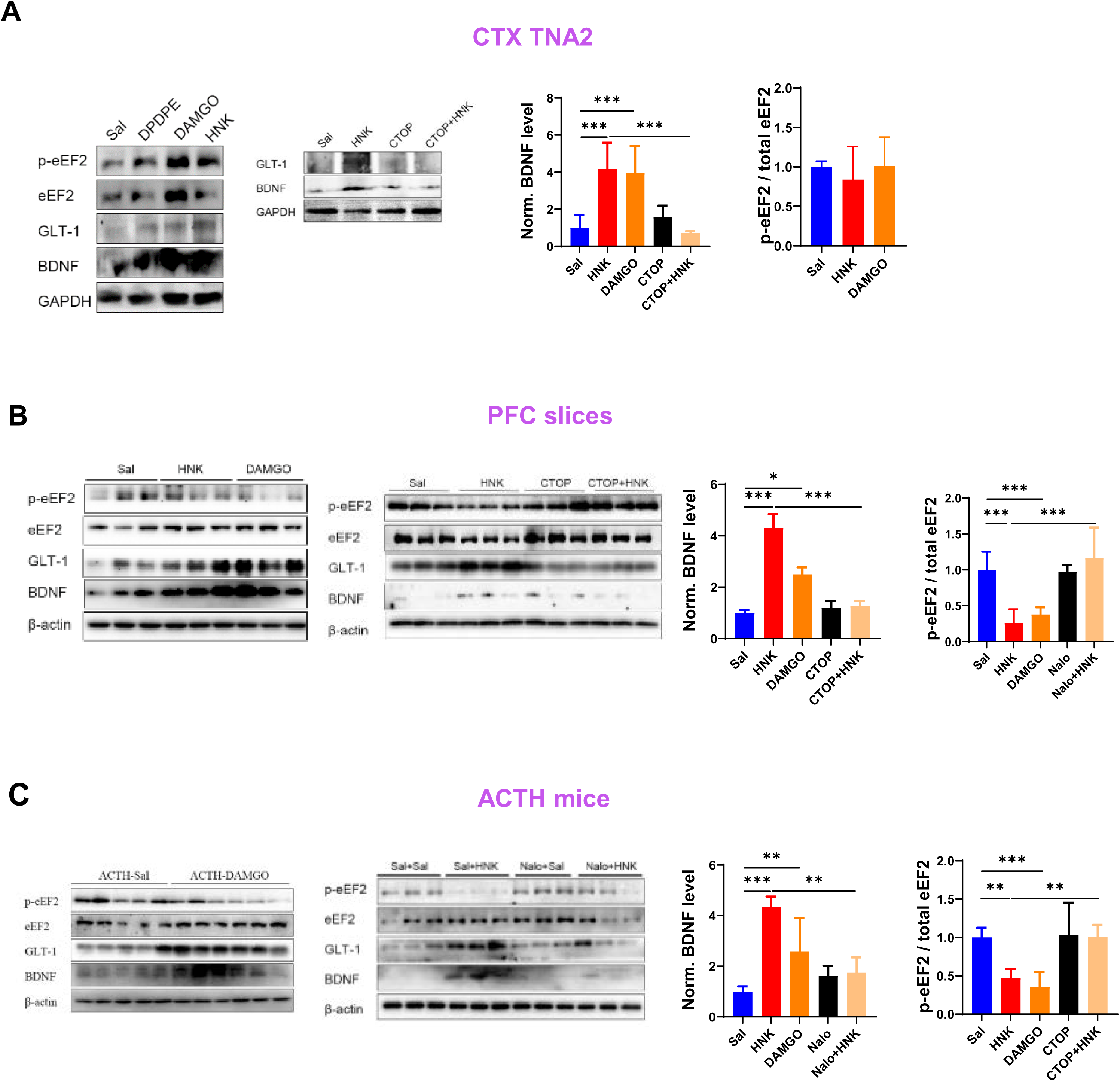
Impact of μ-OR agonists, antagonists on the levels of various proteins. Sample blots (left) and quantification (right) of indicated proteins in CTX TNA2 cells (A; one-way ANOVA; n = 3 experiments), acute PFC slices (B; one-way ANOVA; n = 3 - 6 mice per group), and PFC of ACTH-mice (C; one-way ANOVA; n = 4 - 9 mice per group) treated with μ-OR agonist or antagonists. Data are mean ± s.e.m. *, P< 0.05; **, P< 0.01; ***, P< 0.001.

**Figure S7:**
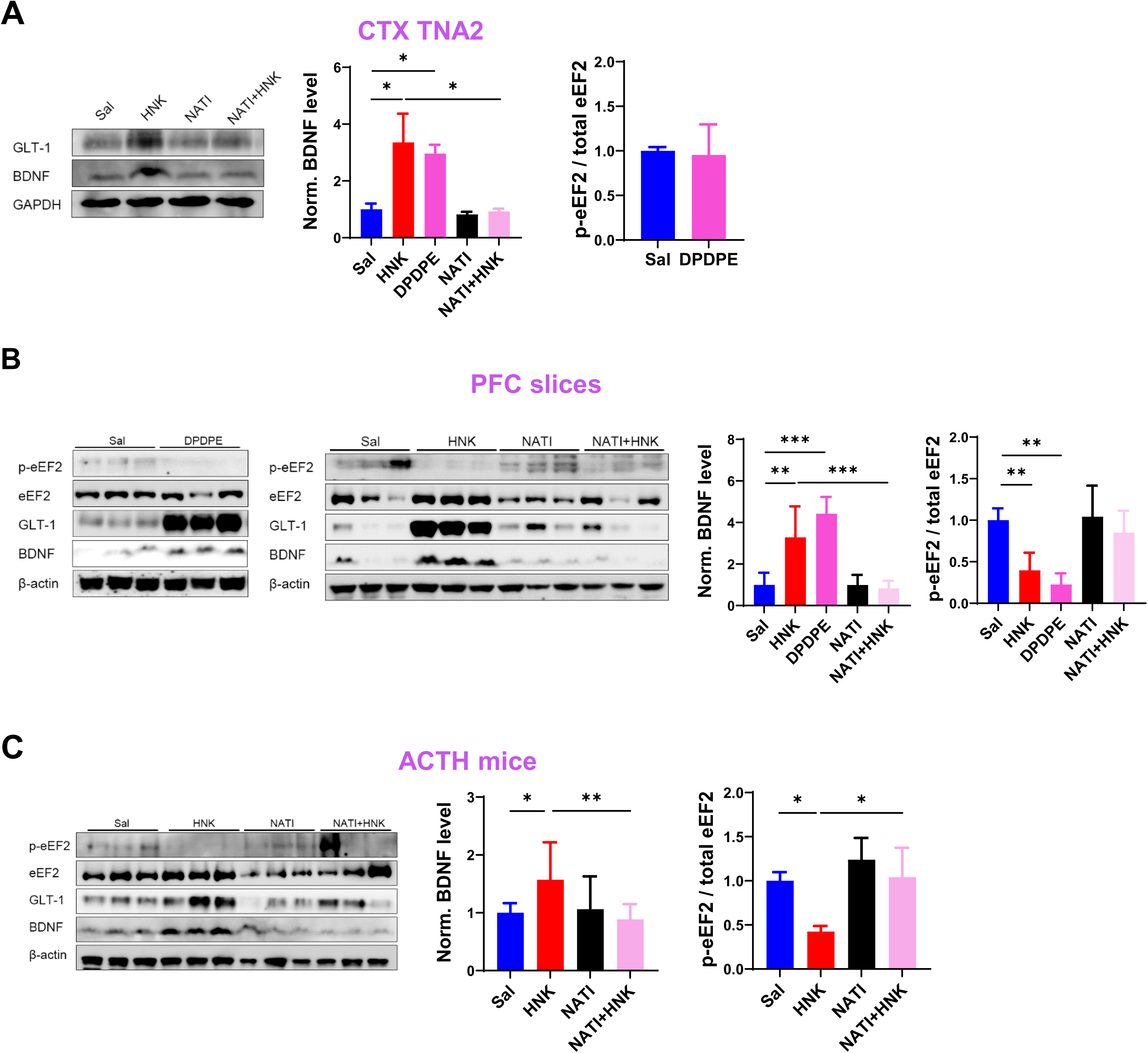
Impact of δ-OR agonists, antagonists on the levels of various proteins. Sample blots (left) and quantification (right) of indicated proteins in CTX TNA2 cells (A; one-way ANOVA; n = 4 experiments), acute PFC slices (B; one-way ANOVA; n = 3 - 6 mice per group), and PFC of ACTH-mice (C; one-way ANOVA; n = 9 - 11 mice per group) treated with δ-OR agonist or antagonist. Data are mean ± s.e.m. *, P< 0.05; **, P< 0.01; ***, P< 0.001.

**Figure S8:**
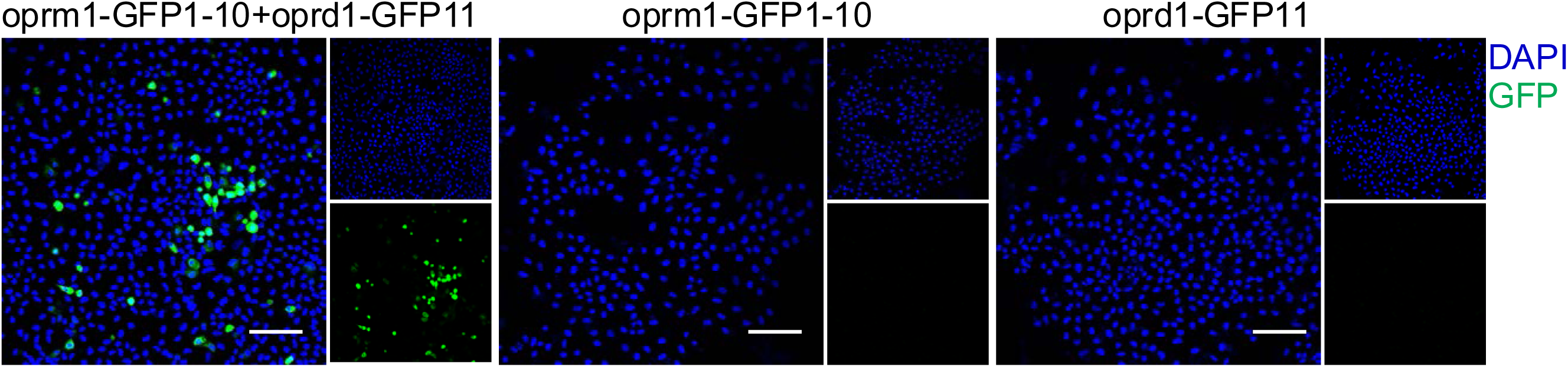
Visualization of μ-δ opioid receptor heterodimerization via split-GFP. Expression of oprm1-GFP1-10, oprd1-GFP11, or co-expression of oprm1-GFP1-10 and oprd1-GFP11, in HeLa cells. Scale bars, 100 μm.

**Figure S9:**
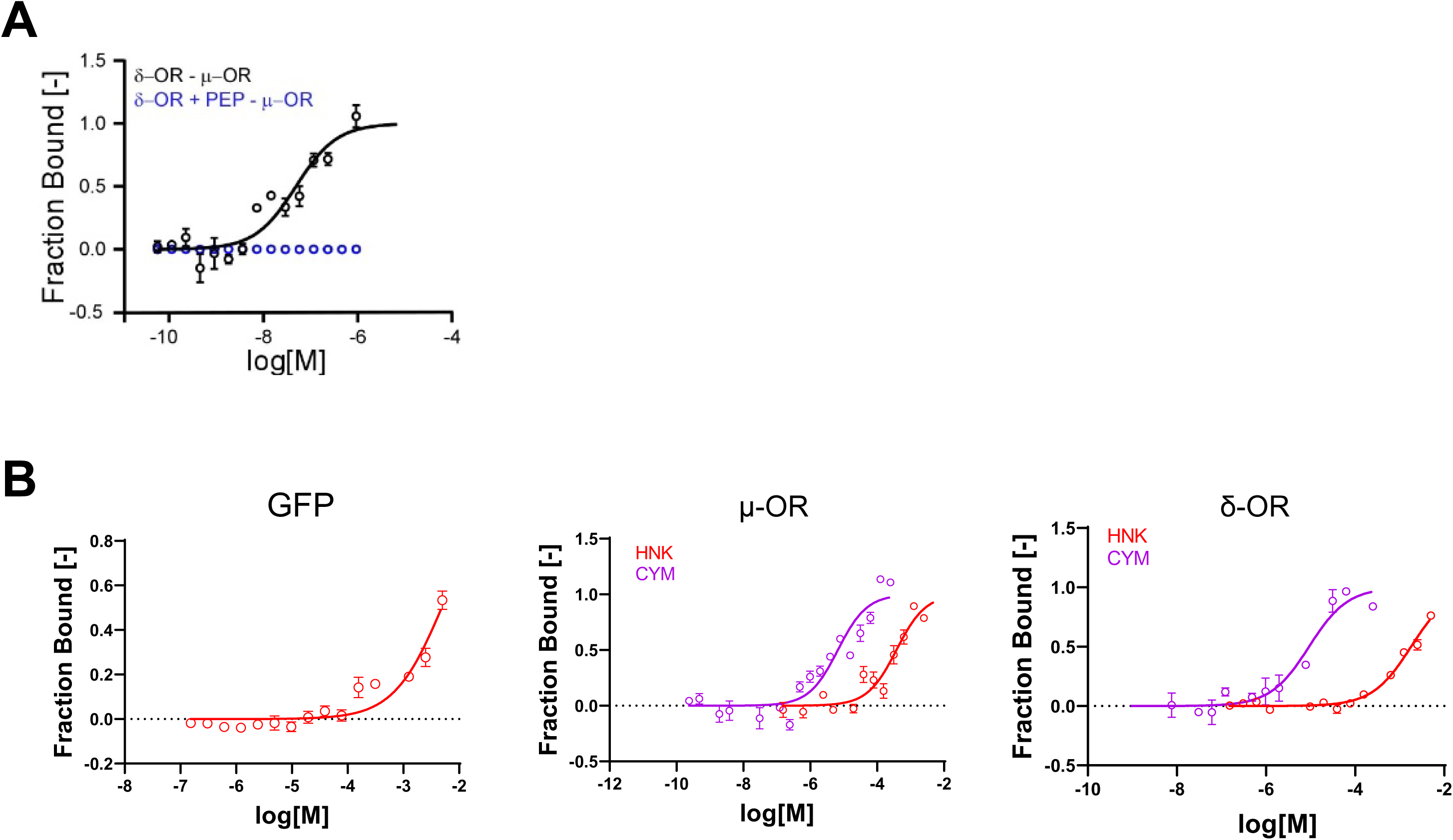
Effects of indicated compounds in lysate from HeLa cells expressing GFP, μ-OR-GFP or δ-OR-mCherry using MST assay. (A) Effect of PEP on the binding between μ-ORs and δ-ORs in MST assay. (B) HNK or CYM MST in lysates from HeLa cells expressing GFP alone, m-OR-ZsGreen, or d-OR-mCherry.

**Figure S10:**
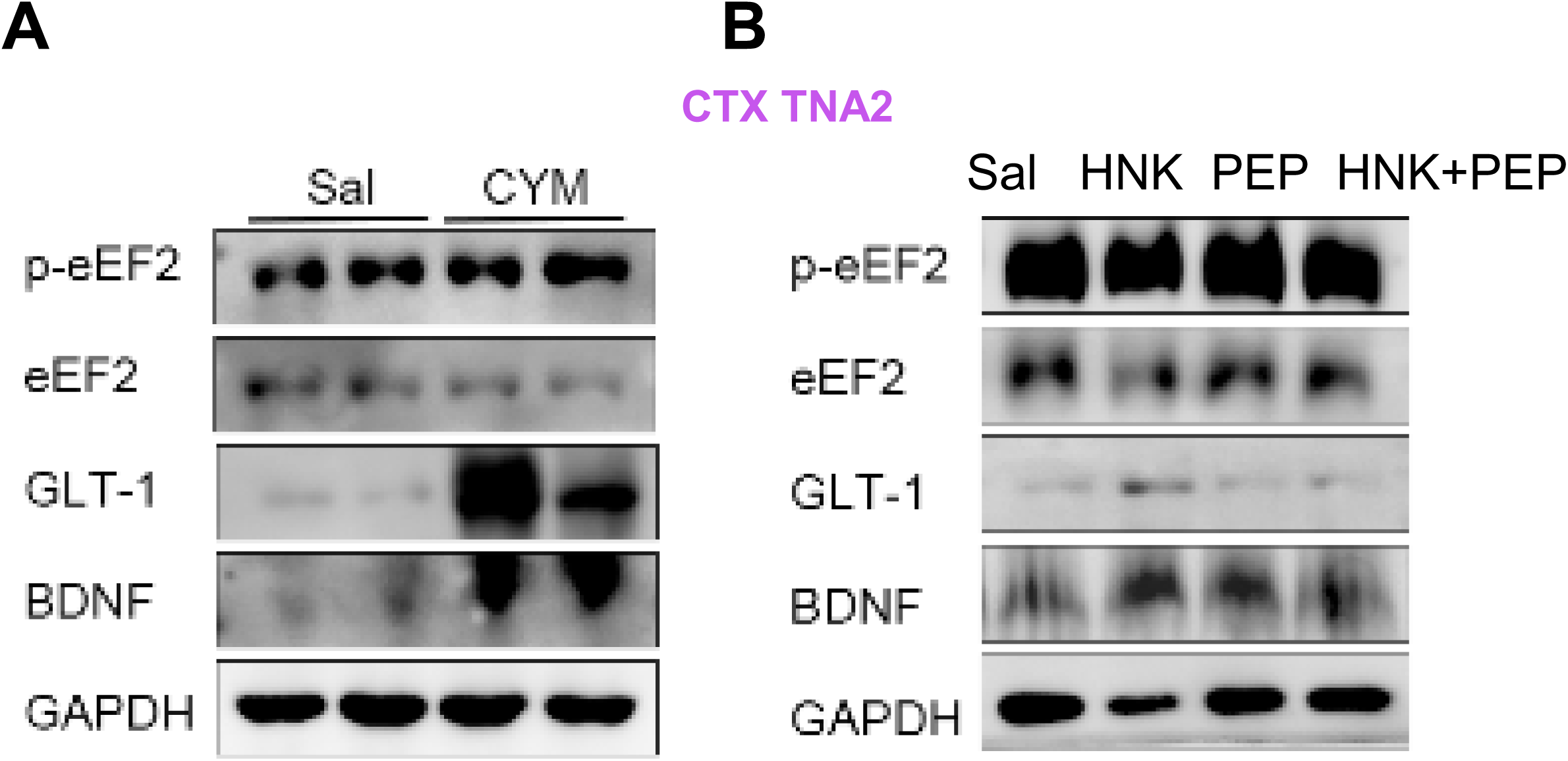
Effect of CYM, HNK and HNK+PEP on GLT-1, BDNF, p-eEF2 /eEF2 in CTX TNA2 cells. Sample image showing the effects of CYM (A) and HNK with or without PEP pre-incubation (B) on GLT-1, BDNF, p-eEF2/eEF2 levels in CTX TNA2 cells.

**Figure S11:**
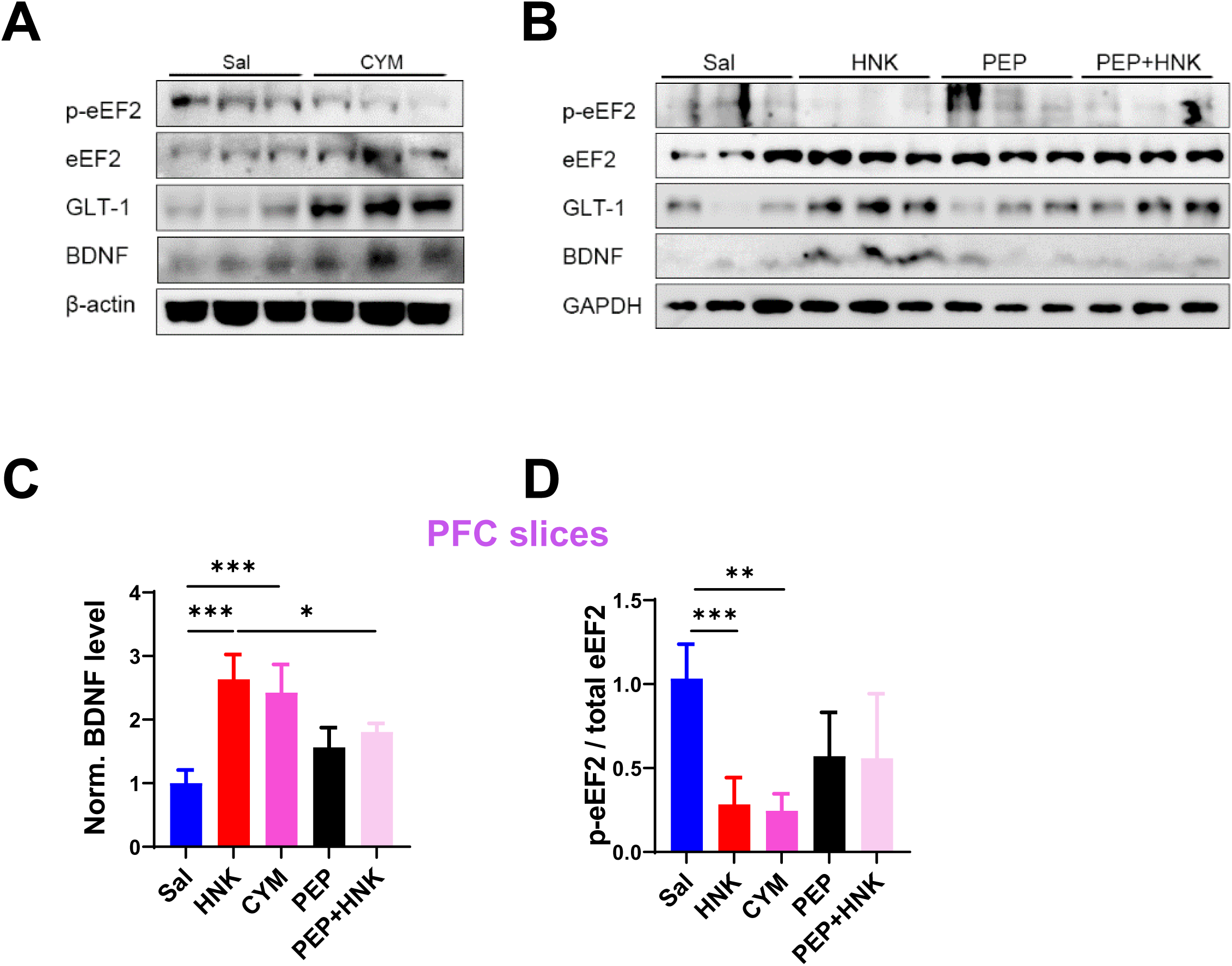
Effect of CYM, HNK and HNK+PEP on GLT-1, BDNF, p-eEF2 /eEF2 in acute PFC slices. (A-B) Sample image of GLT-1, BDNF, p-eEF2 /eEF2 in PFC slices treated with CYM, HNK, HNK+PEP. (C-D) Quantifications on BNDF (C) and p-eEF2 /eEF2 (D; one-way ANOVA; n = 3 - 6 mice per group).

**Figure S12:**
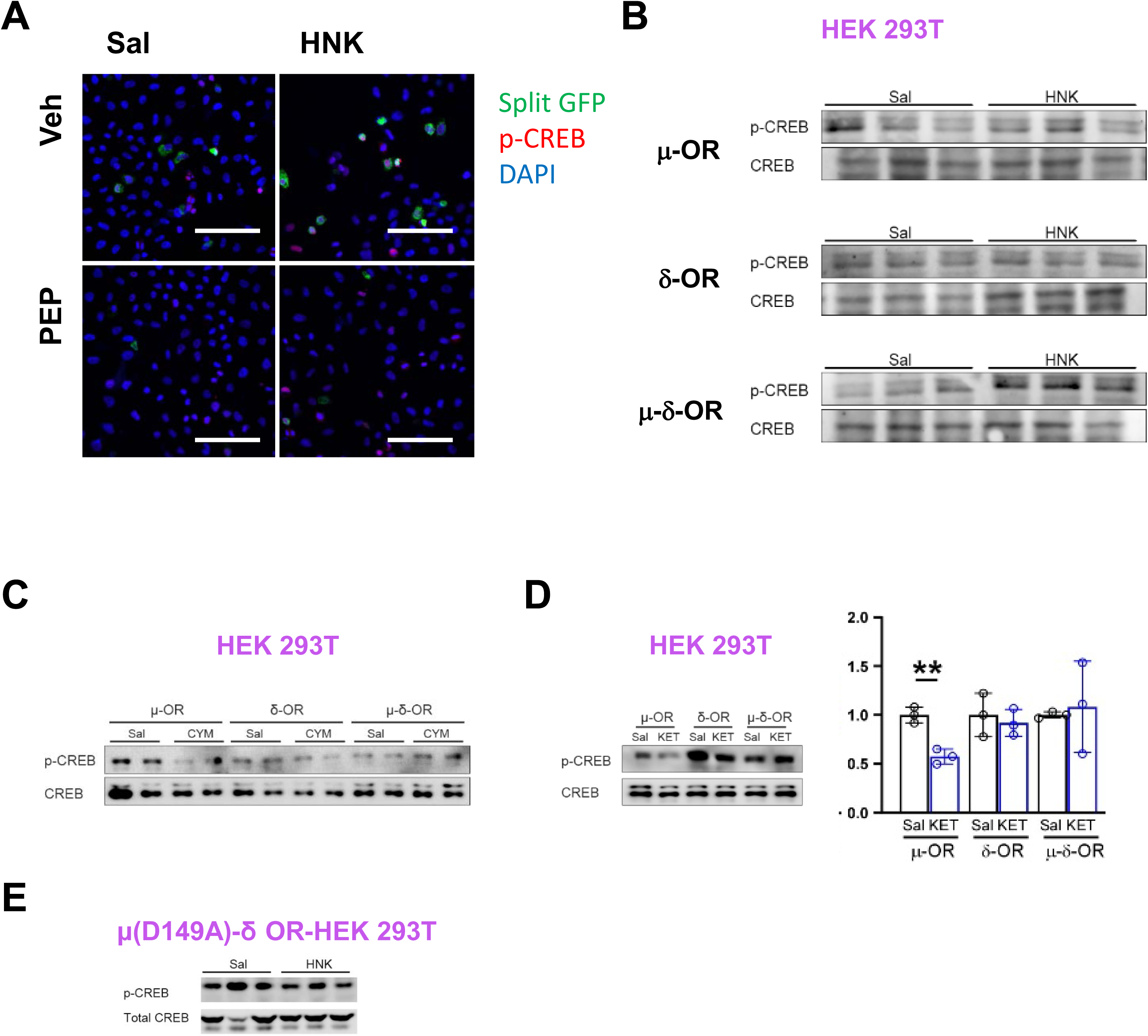
Effect of various compounds on p-CREB/total CREB in HEK 293T cells transfected with μ-ORs, δ-ORs or both μ-ORs and δ-ORs. (A) Sample images showing the effect of PEP pre-incubation on HNK-induced μ-δ-OR-sGFP fluorescence and p-CREB immunostaining in HeLa cells. Scale bar, 100 μm. (B-E) Sample western blot images of p-CREB/CREB ratio in HEK 293T cells expressing the indicated receptors (B, E), CYM (C), or ketamine (D; two-tailed t-test for d; n = 3 experiments). Data are mean ± s.e.m. *, P< 0.05; **, P< 0.01; ***, P< 0.001.

**Figure S13:**
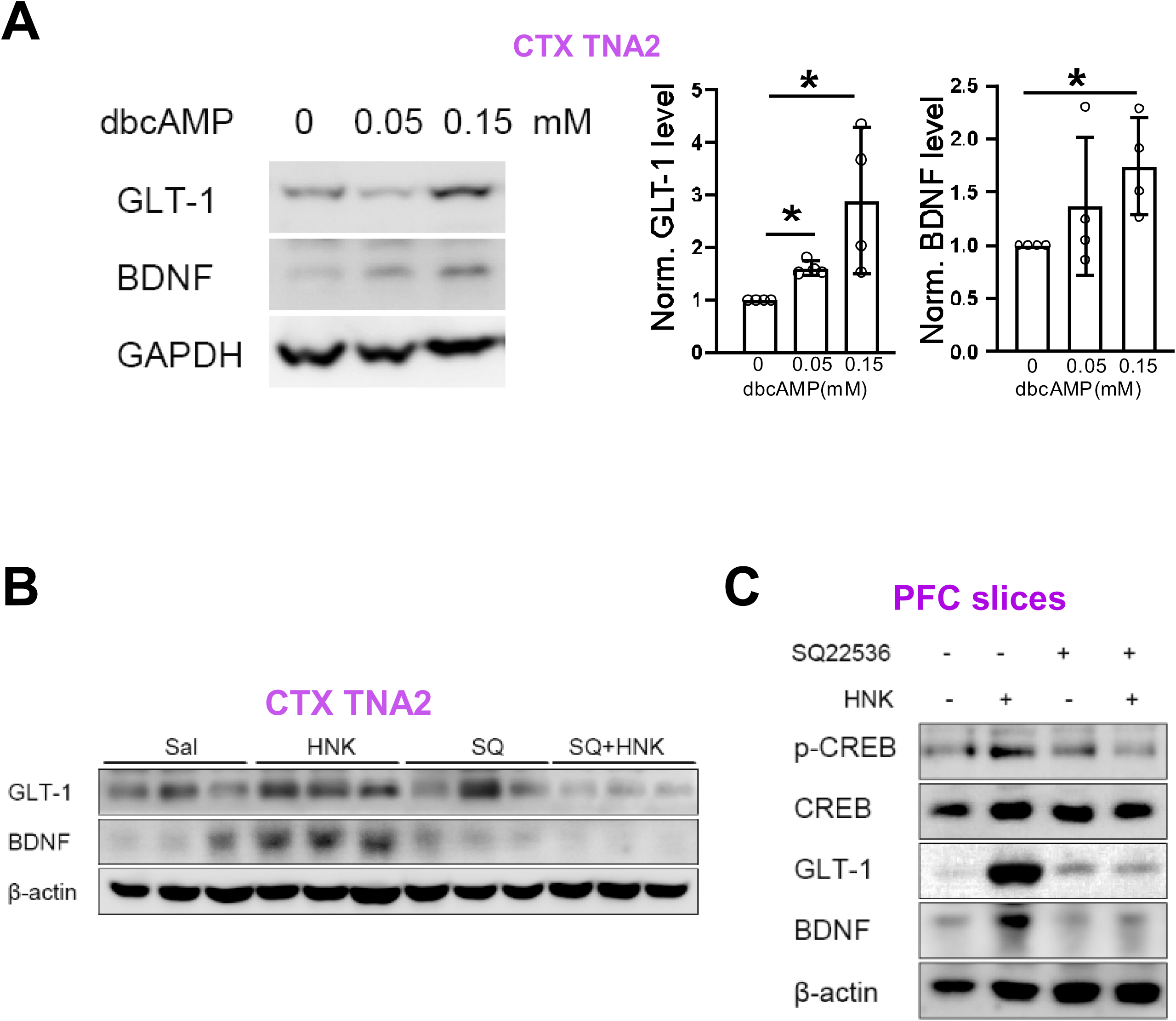
Effect of dbcAMP on levels of GLT-1, BDNF in CTX TNA2 cells. (A) Sample image (left) and quantification of GLT-1 and BDNF levels in CTX TNA2 cells treated with different concentrations of dbcAMP (n = 4 experiments). (B) Sample image showing the effect of adenylate cyclase inhibitor SQ22536 on HNK-induced elevated protein levels of GLT-1, BDNF and p-CREB in CTX TNA2 cells. (C) Sample image showing the effect of adenylate cyclase inhibitor SQ22536 on HNK-induced elevated protein levels of GLT-1, BDNF in PFC slices. Data are mean ± s.e.m. *, P< 0.05; **, P< 0.01; ***, P< 0.001.

**Figure S14:**
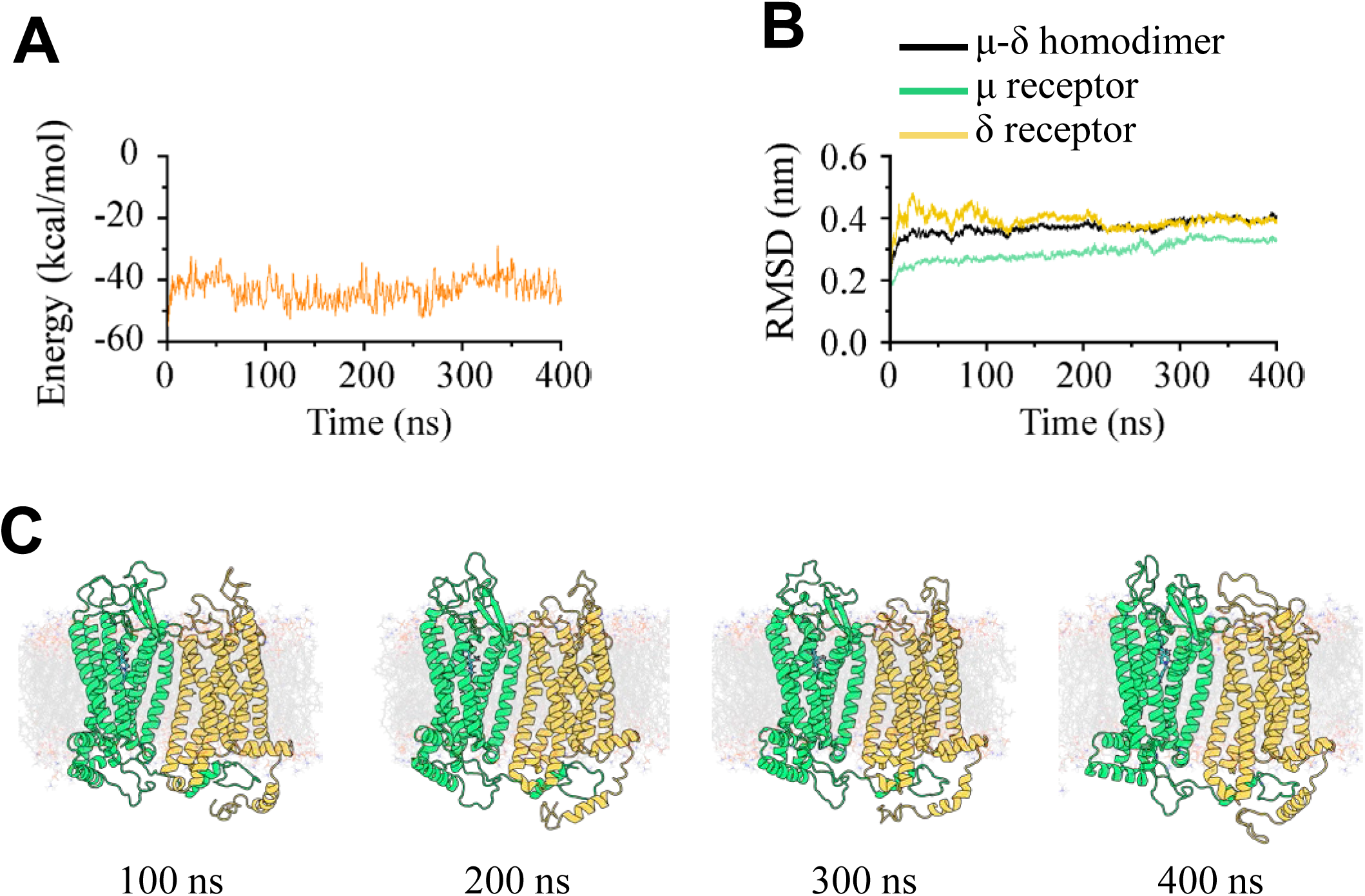
Binding dynamics and equilibrium analysis of CYM with the μ–δ opioid receptor heterodimer. Binding dynamics were assessed by monitoring the time evolution of the binding free energy (A) and the RMSD of backbone atoms (B) for the entire complex as well as for each opioid receptor. Representative structural snapshots are shown every 100 ns (C).

**Figure S15:**
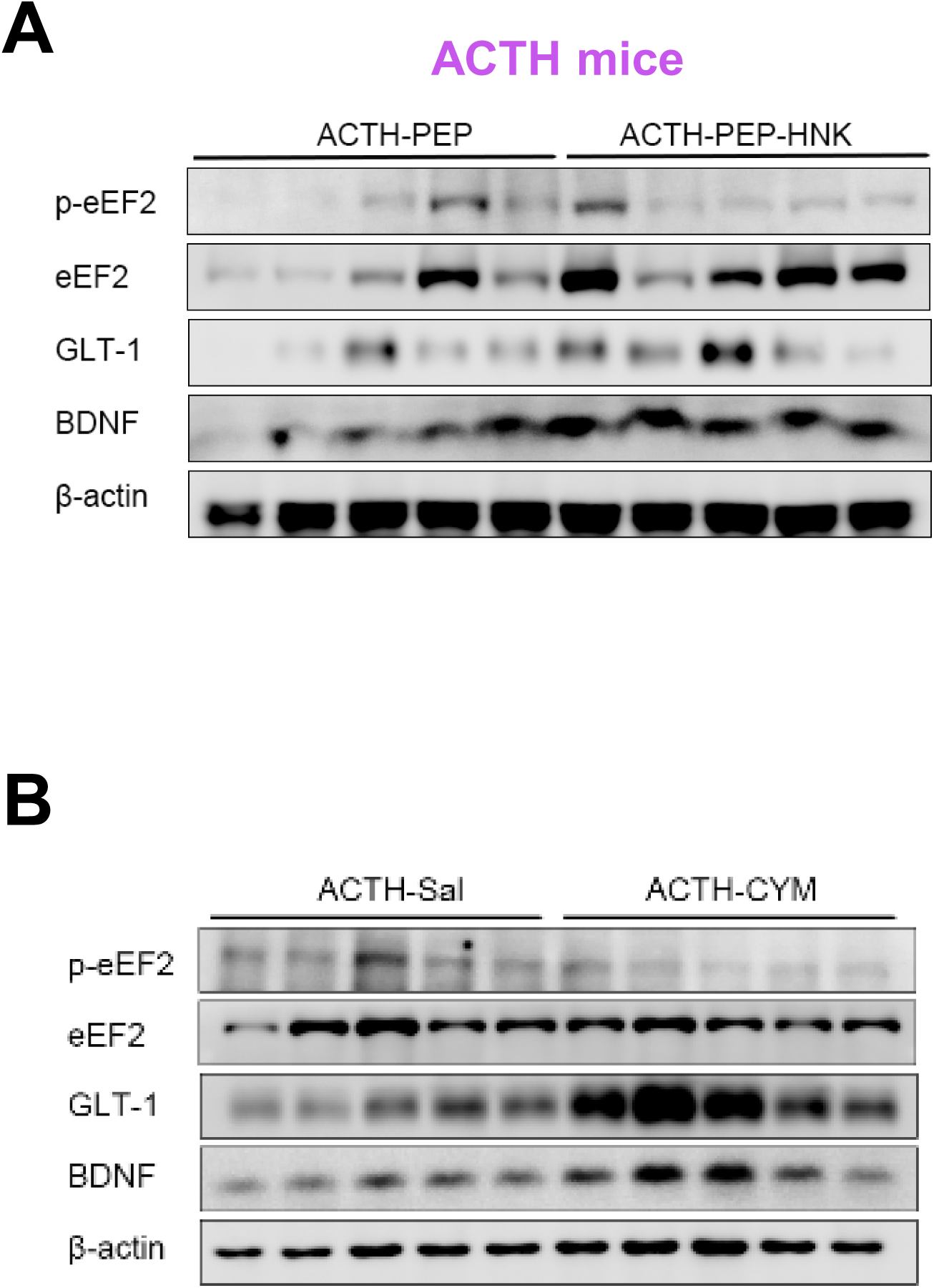
Effect of HNK and HNK+PEP on BDNF, p-eEF2/eEF2 in ACTH mice. (A) Sample blot image of BDNF, p-eEF2/eEF2 in ACTH-mice treated with HNK, HNK+PEP. (B) Sample blot image of BDNF, p-eEF2/eEF2 in ACTH-mice treated with CYM.

